# Real-time denoising of fluorescence time-lapse imaging enables high-sensitivity observations of biological dynamics beyond the shot-noise limit

**DOI:** 10.1101/2022.03.14.484230

**Authors:** Xinyang Li, Yixin Li, Yiliang Zhou, Jiamin Wu, Zhifeng Zhao, Jiaqi Fan, Fei Deng, Zhaofa Wu, Guihua Xiao, Jing He, Yuanlong Zhang, Guoxun Zhang, Xiaowan Hu, Yi Zhang, Hui Qiao, Hao Xie, Yulong Li, Haoqian Wang, Lu Fang, Qionghai Dai

**Affiliations:** Department of Automation, Tsinghua University, Beijing 100084, China; Tsinghua Shenzhen International Graduate School, Tsinghua University, Shenzhen, 518055, China; Institute for Brain and Cognitive Sciences, Tsinghua University, Beijing 100084, China; School of Information and Technology, Fudan University, Shanghai 200433, China; Department of Electronic Engineering, Tsinghua University, Beijing 100084, China; Beijing Key Laboratory of Multi-dimension & Multi-scale Computational Photography (MMCP), Tsinghua University, Beijing 100084, China; IDG/McGovern Institute for Brain Research, Tsinghua University, Beijing, China; State Key Laboratory of Membrane Biology, Peking University School of Life Sciences, Beijing 100871, China; PKU-IDG/McGovern Institute for Brain Research, Beijing 100871, China; Hangzhou Zhuoxi Institute of Brain and Intelligence, Hangzhou, 311100, China

## Abstract

A fundamental challenge in fluorescence microscopy is the inherent photon shot noise caused by the inevitable stochasticity of photon detection. Noise increases measurement uncertainty, degrades image quality, and limits imaging resolution, speed, and sensitivity. To achieve high-sensitivity imaging beyond the shot-noise limit, we provide DeepCAD-RT, a versatile self-supervised method for effective noise suppression of fluorescence time-lapse imaging. We made comprehensive optimizations to reduce its data dependency, processing time, and memory consumption, finally allowing real-time processing on a two-photon microscope. High imaging signal-to-noise ratio (SNR) can be acquired with 10-fold fewer fluorescence photons. Meanwhile, the self-supervised superiority makes it a practical tool in fluorescence microscopy where ground-truth images for training are hard to obtain. We demonstrated the utility of DeepCAD-RT in extensive experiments, including *in vivo* calcium imaging of various model organisms (mouse, zebrafish larva, fruit fly), 3D migration of neutrophils after acute brain injury, and 3D dynamics of cortical ATP (adenosine 5’-triphosphate) release. DeepCAD-RT will facilitate the morphological and functional interrogation of biological dynamics with minimal photon budget.

## Introduction

The proper functioning of living organisms relies on a series of spatiotemporally orchestrated cellular and subcellular activities. Observing and recording these phenomena is considered to be the first step towards understanding them. Fluorescence microscopy, combined with the growing palette of fluorescent indicators, provides biologists with a practical tool capable of good molecular specificity and high spatiotemporal resolution. Recent advances in fluorescence imaging have brought us insights into various previously inaccessible processes, ranging from organelle interactions at nanoscale^1–3^, to pan-cell footprints during embryo development^4–6^, and to whole-brain neuronal dynamics synchronized with certain behaviors^7–10^.

Among the challenges of fluorescence microscopy, poor imaging SNR caused by limited photon budget lingeringly stands in the central position. The causes of this photon-limited challenge are manifold. Firstly, the low photon yield of fluorescent indicators and their low concentration in labeled cells result in the lack of photons at the source. Secondly, although using higher excitation power is a straightforward way to increase fluorescence photons, living systems are too fragile to tolerate high excitation dosage. Extensive experiments have shown that illumination-induced photobleaching, phototoxicity, and tissue heating will disturb crucial cellular processes including cell proliferation, migration, vesicle release, neuronal firing, etc^11–18^. Thirdly, recording fast biological processes necessitates high imaging speed and the short dwell time further exacerbates the shortage of photons. Finally, the quantum nature of photons makes the stochasticity (shot noise) of optical measurements inevitable^19, 20^. The intensity detected by photoelectric sensors follows a Poisson distribution parameterized with the exact photon count^21^. In fluorescence imaging, detection noise dominated by photon shot noise aggravates the measurement uncertainty and obstructs the veritable visualization of underlying structures, potentially altering morphological and functional interpretations that follow. To capture enough photons for satisfactory imaging sensitivity, researchers have to sacrifice imaging speed, resolution, and even sample health^19, 22^.

Comprehensive efforts have been invested to increase the photon budget of fluorescence microscopy, from designing high-performance fluorophores^23–25^, to upgrading the excitation and detection physics^19, 26–28^, and to developing data-driven denoising algorithms^22, 29–31^. We previously developed DeepCAD, a deep self-supervised denoising method for calcium imaging data, which effectively suppresses the detection noise and improves imaging SNR more than 10-fold without requiring any high-SNR observations^32^. A single low-SNR calcium imaging sequence can be directly used as the training data to train a denoising convolutional neural network.

Here, with advancements in methods and applications, we present DeepCAD-RT, a versatile self-supervised denoising method for fluorescence time-lapse imaging with real-time processing speed and improved performance. By pruning redundant features inside the network architecture, we constructed a lightweight network and compressed the model parameters by 94%, which consequently reduced 85% processing time and 70% memory consumption. Meanwhile, we augmented the training data by 12-fold to alleviate the data dependency and make the method still tractable with a small amount of data. We show that such a strategy of combining model compression and data augmentation eliminates overfitting and makes the training process stable and manageable. Finally, we optimized the hardware deployment of DeepCAD-RT and achieved an overall improvement of a 27-fold reduction in memory consumption and a 20-fold acceleration in inference speed, which ultimately supported real-time image denoising once incorporated with the microscope acquisition system. We demonstrate the capability and generality of DeepCAD-RT on a series of photon-limited imaging experiments, including imaging calcium transients in various model organisms such as mice, zebrafish, and flies, observing the migration of neutrophils after acute brain injury, and monitoring cortical neurotransmitter dynamics using a recently developed genetically encoded ATP sensor^33^.

## Results

### Comprehensive optimization of DeepCAD-RT for real-time processing

Limited by the computationally demanding nature of deep neural networks, the throughput of most deep-learning-based methods for video processing is lower than the data acquisition rate^34^. To the best of our knowledge, no deep-learning-based denoising methods for fluorescence imaging have been demonstrated to have real-time processing capability in practice. The original DeepCAD was proposed to denoise calcium imaging data in post-processing. For the same amount of data, its processing time is about five times longer than the acquisition time. Differently, in this work, our rationale was to provide a compact and user-friendly tool that can be incorporated into the data acquisition pipeline to enhance the raw noisy data immediately after acquisition, which serves as the last step of data acquisition and the first step of data processing. Towards this goal, we started the first round of optimization by simplifying the network architecture (Fig. 1a). We compressed the network by pruning different proportions of network parameters and then investigated their performance using synthetic calcium imaging data simulated with NAOMi^35^. Synthetic calcium imaging data have paired ground truth images that are indispensable for rigorous comparison (Supplementary Fig. 1). Quantitative evaluation shows that although we removed as many as ~94% (from 16.3 million to 1.0 million) network parameters, the denoising performance did not deteriorate (Supplementary Fig. 2) while the memory cost and inference time were reduced respectively by 3.3-fold and 6.6-fold, which pushed the processing throughput of the network to the same level as imaging (Fig. 1b). However, unlike denoising in post-processing, real-time processing requires frequent data exchanges and necessitates extra computational resources for display and interaction. A practical processing throughput should be 2-3 times higher than imaging to reserve reasonable design margins. For further acceleration, we carried out the second round of optimization in hardware deployment by implementing simplified models with TensorRT (NVIDIA), a toolbox that provides optimized deployment of deep neural networks on specific graphics processing unit (GPU) cards. On our task, the deployment optimization reduced the memory cost and inference time by 8.2-fold and 3.0-fold, respectively. Combining model simplification and deployment optimization, the overall improvement is a 27-fold reduction in memory consumption and a 20-fold improvement in inference speed (Fig. 1b), making the implementation of real-time denoising possible.

**Fig. 1.**
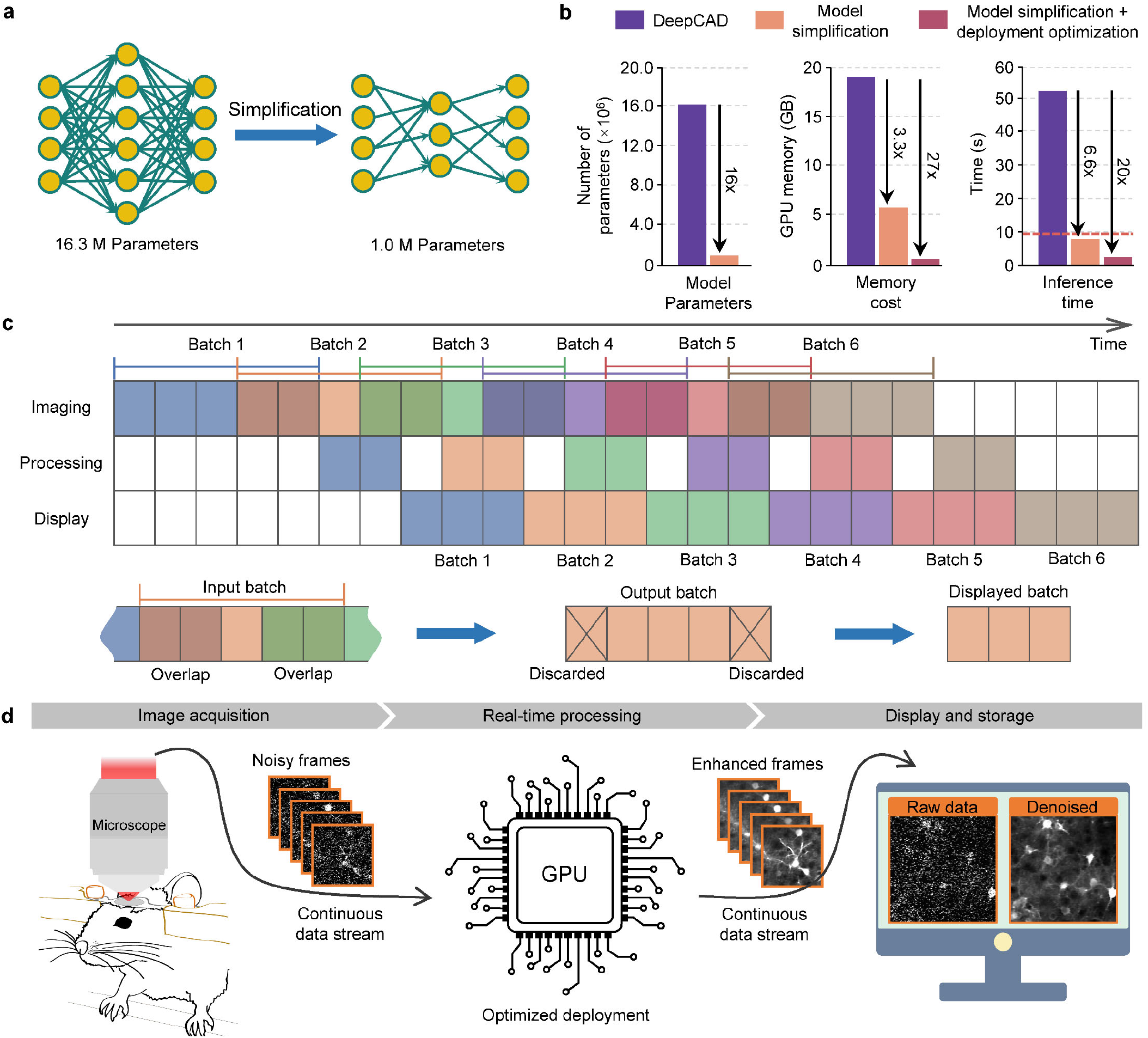
Optimization and real-time schedule of DeepCAD-RT. **a**, Model simplification by feature pruning. The total number of model parameters was reduced from ~16.3 million (16,315,585) to ~1.0 million (1,020,337) for higher processing speed and less memory consumption. **b**, Performance comparison between DeepCAD and DeepCAD-RT. Deployment optimization refers to hardware acceleration by further optimizing the deployment of deep neural networks on graphics processing unit (GPU) cards. An example image sequence of 490×490×300 (x-y-t) pixels was partitioned into 75 patches (150×150×150 pixels 40% overlap) to obtain these performance measurements on the same GPU (GeForce RTX 3090, Nvidia) with one batch size. Totally, ~2.53×10^8^ pixels flowed through the network. All hyperparameters remained the same except the method. The red dashed line in the rightmost panel indicates the imaging time (~9.6 s) of the example data. **c**, Real-time schedule of DeepCAD-RT. Continuous data stream acquired from the microscope acquisition software was packaged into 3D (x-y-t) mini-batches and then fed into DeepCAD-RT. To maximize the processing speed, three parallel threads were programmed for image acquisition, data processing, and display, respectively. For each batch, half of the overlap was discarded to avoid marginal artifacts. Overlapping frames between two consecutive batches are rendered with overlapping colors. **d**, Schematic of real-time denoising implemented with DeepCAD-RT on a two-photon microscope. Raw noisy data and the corresponding denoised data are displayed synchronously, which will be saved as separated files automatically at the end of the imaging session.

To incorporate DeepCAD-RT into the data acquisition pipeline of the microscopy system, we designed three parallel threads for imaging, data processing, and display (Fig. 1c). The continuous data stream captured by the microscope will be packaged into consecutive batches in the imaging thread and then seamlessly fed into the processing thread. Once a new batch is received by the processing thread, the pre-trained model already deployed on GPU starts processing and the denoised batch will be passed to the display thread. After removing overlapping frames, denoised batches will be assembled into a denoised stream and displayed on the monitor. The three threads keep temporally aligned throughout the whole imaging session. Both the raw noisy data and denoised data will be saved as separated files once the imaging session finishes. As a proof-of-concept, we demonstrate real-time denoising on a two-photon fluorescence microscope using DeepCAD-RT (Fig. 1d and Supplementary Fig. 3). The denoised data with drastically enhanced SNR can be presented simultaneously with the raw data (Supplementary Video 1), which facilitates the observation and evaluation of biological dynamics in photon-limited conditions.

Besides real-time denoising, we also optimized the training procedure to make DeepCAD-RT easy to harness in various biological applications. We introduced 12-fold data augmentation (Supplementary Fig. 4) to reduce its data dependency. Currently, training the network with a low-SNR video stack containing as few as 1000 frames is sufficient to ensure satisfactory performance (Supplementary Fig. 5). Moreover, we found that the combination of model simplification and data augmentation eliminates overfitting (Supplementary Fig. 6), which was an inherent problem of self-supervised training and required human inspections for model selection previously^32^. We compared DeepCAD-RT with DeepInterpolation, another recently developed denoising method leveraging inter-frame correlations^31^. The results show that, with the same amount of training data, DeepCAD-RT significantly outperformed DeepInterpolation, especially in photon-limited conditions (SNR < 5 dB). On the other side, DeepCAD-RT can achieve comparable performance with tens of times less training data (trained from scratch with 6000 frames) than DeepInterpolation (pre-trained with 225,000 frames and then fine-tuned with 6000 frames) (Supplementary Fig. 7). The high data efficiency of DeepCAD-RT enables it to be extended to other applications beyond calcium imaging (Supplementary Fig. 8). In most cases, the data at hand can be directly used for training without requiring additional large-scale training datasets. Another advantage of DeepCAD-RT is that its processing speed can be at least an order of magnitude higher than DeepInterpolation even with the same network complexity and device since DeepCAD-RT outputs the entire 3D stack from the 3D input while DeepInterpolation just outputs a single frame from the 3D input.

### Denoising calcium imaging on multiple model organisms

Although synthetic data can provide ground-truth images that are not experimentally available, the performance of denoising methods should be quantitively evaluated with experimentally obtained data for best reliability. Motivated by this principle, we captured synchronized low-SNR and high-SNR image pairs with our custom-designed two-photon microscope (Supplementary Fig. 9) for each type of experiment. The low-SNR data were used as the input while the synchronized high-SNR data with 10-fold SNR were used for result validation (Supplementary Fig. 10). A standard two-photon microscope was also integrated into our system for cross-system validation and multi-color imaging.

To demonstrate the capability and generality of our method, we first investigated whether it could be applied to various calcium imaging experiments. We began by imaging calcium transient of postsynaptic dendritic spines in cortical layer 1 (L1) of a mouse expressing genetically encoded GCaMP6f^36^. Technically, calcium imaging of dendritic spines over a large field-of-views (FOV) is particularly challenging because of their small sizes^37^. Each spine is usually characterized by as few as several pixels and noise severely contaminates its spatiotemporal features. After we enhanced the original low-SNR data with our method, the image SNR was substantially improved and postsynaptic structures can be clearly resolved even in a single frame (Fig. 2a and Supplementary Video 2). Without noise contamination, the morphological heterogeneity between mushroom spines and stubby spines became discernable. Since different spine classes have different functions during development and learning^38^, revealing spine morphology is helpful for the study of dendritic computing. For quantitative evaluation, we extracted image slices along three dimensions (x-y-t) and calculated image correlations with corresponding high-SNR images. Statistical analysis shows that image correlations can be significantly improved for all three dimensions after denoising (Fig. 2b), manifesting the spatial and temporal denoising capability of our method.

**Fig. 2.**
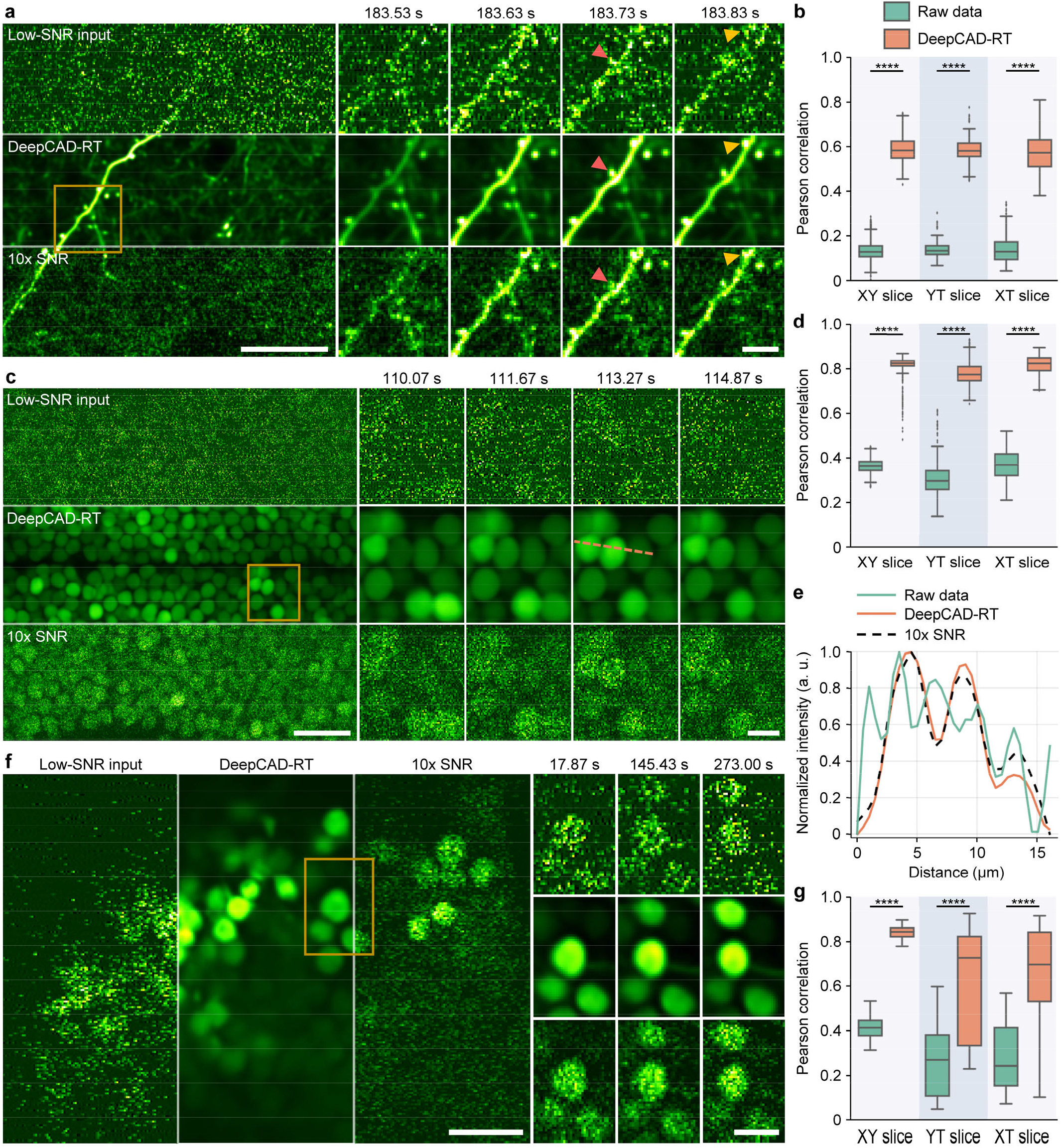
Universal denoising of calcium imaging in mouse, zebrafish, and *Drosophila*. **a**, Imaging calcium transients in dendritic spines of a mouse expressing genetically encoded GCaMP6f calcium indicator. One example frame is shown for the low-SNR raw recording (top), DeepCAD-RT denoised recording (middle), and synchronized high-SNR recording with 10-fold SNR (bottom). Magnified views of the yellow boxed region show calcium dynamics of two spatially adjacent dendritic branches. Each frame was integrated for 33 ms to ensure high temporal resolution. Red arrowheads point to a mushroom spine and yellow arrowheads point to a stubby spine. Scale bar, 20 μm for the whole fìeld-of-view (FOV) and 5 μm for magnified views. **b**, Boxplots showing image correlations along three dimensions (x-y-t) before and after denoising. The high-SNR data with 10-fold SNR was used as the reference for correlation computing. XY slice, N=6000; YT slice, N=246, XT slice, N=489. **c**, Time-lapse imaging of calcium dynamics of optic tectum neurons in the zebrafish brain (HuC:GCaMP6s). Top, the original low-SNR data. Middle, DeepCAD-RT enhanced data. Bottom, high-SNR recording with 10-fold SNR. Magnified views show the neural activity of the yellow boxed region in a short period. Each frame was integrated for 66 ms. Scale bar, 20 μm for the entire FOV and 5 μm for magnified views. **d**, Pearson correlations of image slices along three dimensions before and after denoising. XY slice, N=6000; YT slice, N=246, XT slice, N=246. **e**, Intensity profiles of the yellow dashed line in **c.** Pixels intensities were extracted from 2-fold down-sampled images and all traces were smoothed by moving average with a 3-pixel kernel to suppress the noise. **f**, Denoising performance of DeepCAD-RT on calcium imaging of *Drosophila* mushroom body (GCaMP7f). The same frame is shown for the original low-SNR data (left), DeepCAD-RT denoised image (middle), and high-SNR image with 10-fold SNR (right). Magnified views show snapshots of the yellow boxed region at three moments. Each frame was integrated for 33 ms. Scale bar, 10 μm for the whole FOV and 5 μm for magnified views. **g**, Boxplots showing the improvement of image correlation after denoising. XY slice, N=12000; YT slice, N=241, XT slice, N=335. Asterisks denote significance levels tested with one-sided paired t-test. ****P < 0.0001 for all comparisons.

Animal models currently used in systems and evolutionary neuroscience are diverse that extend from jellyfish^39^ to monkeys^40^. To test our method on versatile animal models with different neuron morphologies and brain structures, we imaged *in vivo* calcium dynamics in the brain of zebrafish larvae and *Drosophila* and then denoised the original shot-noise-limited signals with our method. For zebrafish imaging, we used larval zebrafish expressing nuclear-localized GCaMP6s calcium indicator throughout the whole brain. Because of the shot noise, raw images deteriorated severely and neurons can be barely recognized. However, after denoising, the image SNR was improved more than 10-fold and fluorescence signals became clear (Fig. 2c and Supplementary Video 3). Image correlations along all three dimensions were significantly improved (Fig. 2d). In each frame, the distribution of optic tectum neurons can be clearly recognized with the enhancement of our method (Fig. 2e). Additionally, we also imaged calcium events of large neuronal populations spanning multiple brain regions and found that the removal of noise was rather helpful for separating densely labeled cells. (Supplementary Fig. 11 and Supplementary Video 4). Similarly, we performed time-lapse calcium imaging of mushroom body neurons in the brain of adult *Drosophila*. The results show that the enhanced imaging SNR and image correlations could facilitate the observation of calcium dynamics (Fig. 2f,g and Supplementary Video 5), which verified the effectiveness of our method on various calcium imaging applications involving different animal models and neuronal structures. Since smaller animals such as zebrafish and *Drosophila* are less resistant to high excitation power than mice, it is difficult to keep the sample healthy and obtain high-SNR imaging data simultaneously. With its good performance and versatility, DeepCAD-RT can be a promising tool for calcium imaging to minimize the excitation power and photon-induced disturbance by removing the shot noise computationally.

### Observing neutrophil migration *in vivo* with low excitation power

Our previous work only focused on calcium imaging, in which neurons are spatially invariant and their intensity changes over time. Next, we applied our method to the observation of cell migration, a complementary task with almost temporally invariant intensity and continuously changing cell positions. Neutrophils are the most abundant white blood cells in immune defense^41^. To fully understand the function of neutrophils, intravital imaging with minimal illumination is essential because phototoxicity and photodamage would alter cellular and subcellular processes, which potentially disturb normal immune response^15, 42^. We first evaluated the performance of our method on cell migration observations qualitatively and quantitatively with synchronized low-SNR and high-SNR (10-fold SNR) image pairs captured by our customized system. The results show that DeepCAD-RT can restore neutrophils of different shapes from noise, as well as the evolution of morphological features over time (Fig. 3a-c and Supplementary Video 6). Since the SNR of denoised data is better than high-SNR data of 10-fold SNR, the illumination power can be equivalently reduced more than 10-fold for linear microscopy and more than 3-fold for two-photon microscopy. For better comparison, we show the kymographs (x-t projections) of marked regions. The migration of neutrophils could be visualized directly in denoised data rather than submersed in noise in low-SNR raw data (Fig. 3d). Quantitative evaluation also indicates that denoised data are more correlated to high-SNR data (Fig. 3e). Additionally, the more than 10-fold improvement in image SNR after denoising prompted us to investigate whether our method could reveal more cellular traits if it took high-SNR data as the input. After training and inference with the high-SNR data, we found that higher input SNR could produce much better denoising results. The dynamics of reaction fibers during neutrophil migration could be visualized after the enhancement of our method (Fig. 3f and Supplementary Video 7).

**Fig. 3.**
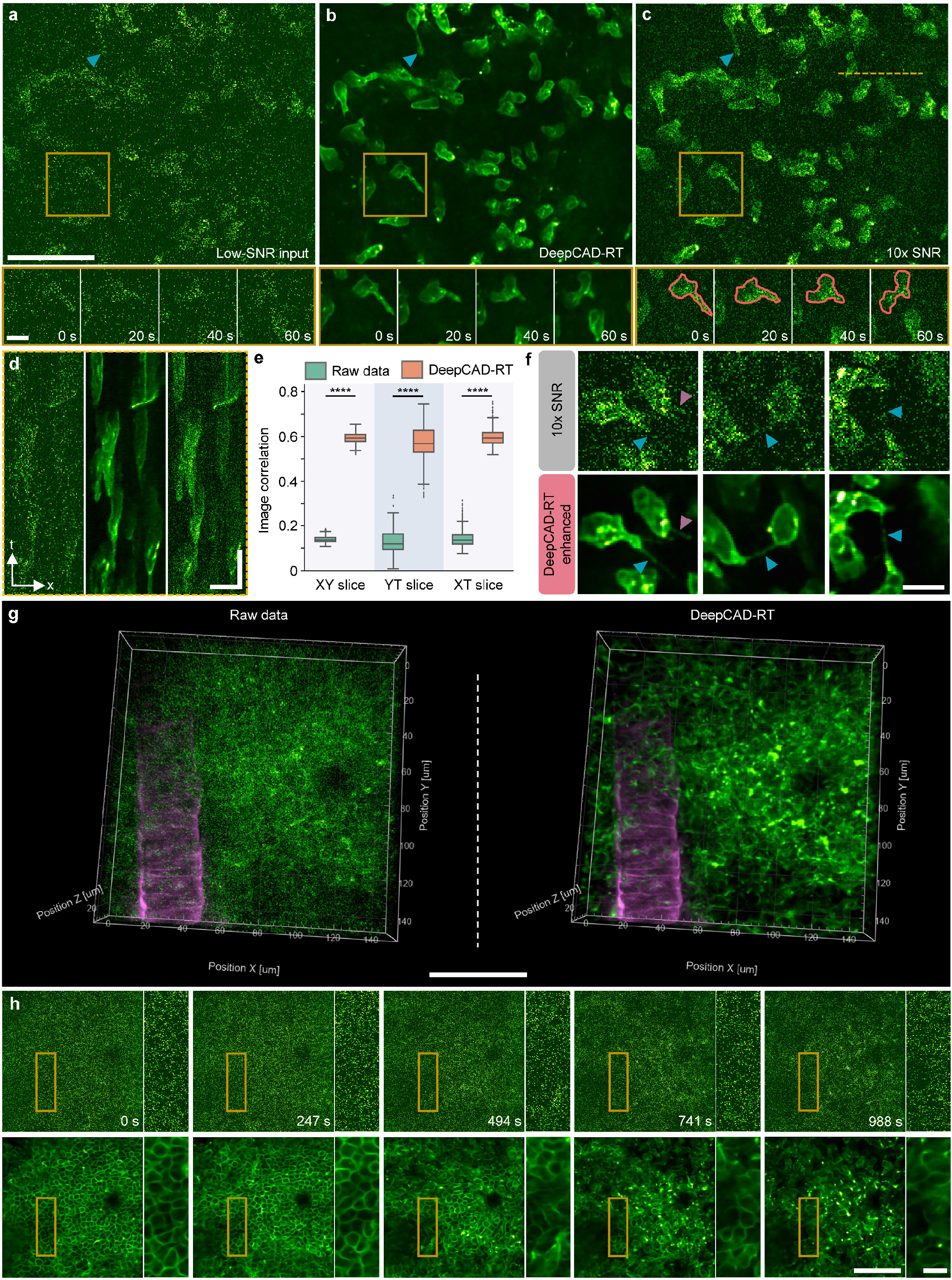
Observing 3D migrations of neutrophils in the mouse brain *in vivo*. **a**, Low-SNR images of neutrophil migration without denoising. **b**, Images denoised with DeepCAD-RT. **c**, Synchronized high-SNR images with 10-fold SNR. Blue arrowheads point to the elongated tail of a migrating neutrophil. Magnified views of the yellow boxed region showing the morphological evolution of neutrophils in a 60 s time window. Red closed lines annotate the border of a neutrophil during migration. Neutrophils were labeled with a fluorescent-conjugated Ly-6G antibody. Each frame was integrated for 100 ms and the entire time-lapse imaging session lasted 644 s. Scale bar, 50 μm for the whole FOV and 10 μm for magnified views. **d**, XT slices along the yellow dashed line in **c** of low-SNR raw data (left), DeepCAD-RT denoised data (middle), and corresponding high-SNR data with 10-fold SNR (right). Scale bar, 20 μm for x and 50 s for t. **e**, Boxplots showing Pearson correlations of image slices along three dimensions (x-y-t) before and after denoising. XY slice, N=6440; YT slice, N=512, XT slice, N=512. P values were calculated by one-sided paired t-test. ****P < 0.0001 for all comparisons. **f**, Denoising high-SNR data with DeepCAD-RT reveals subcellular dynamics of neutrophils. Reaction fibers are indicated with arrowheads. Scale bar, 10 μm. **g**, 3D imaging of neutrophil migration in a 150×150×30 μm^3^ volume (15 planes) after acute brain injury. The raw noisy volume (left) and corresponding denoised volume (right) are visualized with the same perspective. Acute brain injury was induced by craniotomy. Neutrophils were labeled with a fluorescent-conjugated Ly-6G antibody (the green channel). Blood vessels were stained with a wheat germ agglutinin (WGA, the magenta channel) dye. Since blood vessels are stationary, noise in the magenta channel was removed by averaging multiple frames. Scale bar, 50 μm. **h**, Images of a single plane before (top) and after (bottom) denoising. DeepCAD reveals the diffusion of the neutrophil population. Magnified views of yellow boxed regions are shown next to each image. Scale bar, 50 μm for the entire FOV and 10 μm for magnified views.

For fluorescence microscopy, denoising is the first step of subsequent data processing and downstream biological analysis. A good denoising method can facilitate cell segmentation, localization, and classification, which are fundamental steps for the study of cell migration. To figure out the improvement our method brings to segmentation, we segmented neutrophils from the original noisy images (both low-SNR and high-SNR) and corresponding denoised images using Cellpose^43^ and Stardist^44^, two recently published methods for cellular segmentation with state-of-the-art performance^45^. We enlisted five expert human annotators to manually label cell borders and obtain ground-truth masks through majority voting (Methods). Using Intersection-over-Union (IoU) score as the metric, the segmentation performance of the two methods could be improved by ~30-fold for low-SNR images (Supplementary Fig. 12). For high-SNR images with 10-fold SNR, we also observed a significant improvement for both methods because shot noise was removed and cell structures could be well recognized after denoising.

The migration of neutrophils is coordinated in 3D. Deciphering its spatiotemporal pattern necessitates volumetric imaging. Using our multi-color two-photon microscope, we imaged a 150×150×30 μm^3^ volume in the mouse brain after acute brain injury induced by craniotomy. The volume rate of the entire imaging session was 2 Hz. Fluorescence signals from neutrophils and blood vessels were recorded simultaneously and then merged into multi-color images *post hoc*. To minimize the interference caused by the excitation laser and record the native pattern of neutrophil migration, the excitation power we used was below 30 mW. Since the fluorescence labeling of neutrophils was only localized to their membranes, the concentration of the fluorophore was low. The SNR of the raw data was very low and cell structures and dynamics could not be visualized because of the contamination of shot noise (Fig. 3g). After we denoised these low-SNR raw data with our method, shot noise can be effectively suppressed and the 3D dynamics of neutrophil migration became explicit (Supplementary Video 8), which unveiled the phenomenon that a cluster of neutrophils congregating in the early stage of inflammation diffused over time (Fig. 3h).

### DeepCAD-RT facilitates the recording of neurotransmitter dynamics

With the recent proliferation of different fluorescent indicators, combining fluorescence microscopy and genetically encoded fluorescent indicators has become a widespread methodology for interrogating the structural, functional, and metabolic mechanisms of living organisms^46^. For the nervous system alone, available activity indicators have gone beyond calcium and already extended to other intracellular and extracellular neurotransmitters including dopamine^47, 48^, GABA (γ-aminobutyric acid)^49^, glutamate^50, 51^, acetylcholine^25, 52^, *etc*. Similar to calcium imaging, shot noise is also a restriction for the imaging of other activity sensors, which reduces the image SNR and limits the *in vivo* characterization and applications of them. To investigate whether our method can be extended to neurotransmitter sensors, we took ATP sensor as an example and recorded cortical ATP release using mice expressing GRAB_ATP1.0_^33^, a recently developed genetically encoded sensor for measuring extracellular ATP (Methods). In the low-SNR raw data, shot noise swamped ATP signals (Fig. 4a). After denoising with our method, these release events can be clearly visualized (Fig. 4b,c and Supplementary Video 9). Kymographs (y-t projections) show that some subtle ATP-release events that could be omitted in the raw data become visible (Fig. 4d-f). Quantitatively, we used corresponding high-SNR images as the ground truth to calculate image correlations along all three dimensions and found that image correlations could be significantly improved after denoising (Fig. 4g). To compare ATP traces before and after denoising, we manually annotated 80 firing sites from the heatmap of peak ΔF/F_0_ (Fig. 4h) and then extracted fluorescence traces representing ATP activity over time. We calculated Pearson correlations between all traces and the ground truth (traces extracted from the high-SNR data). Statistical results show that the signals of ATP release can be effectively enhanced and the correlations of all fluorescence traces were improved benefiting from the removal of noise (Fig. 4i).

**Fig. 4.**
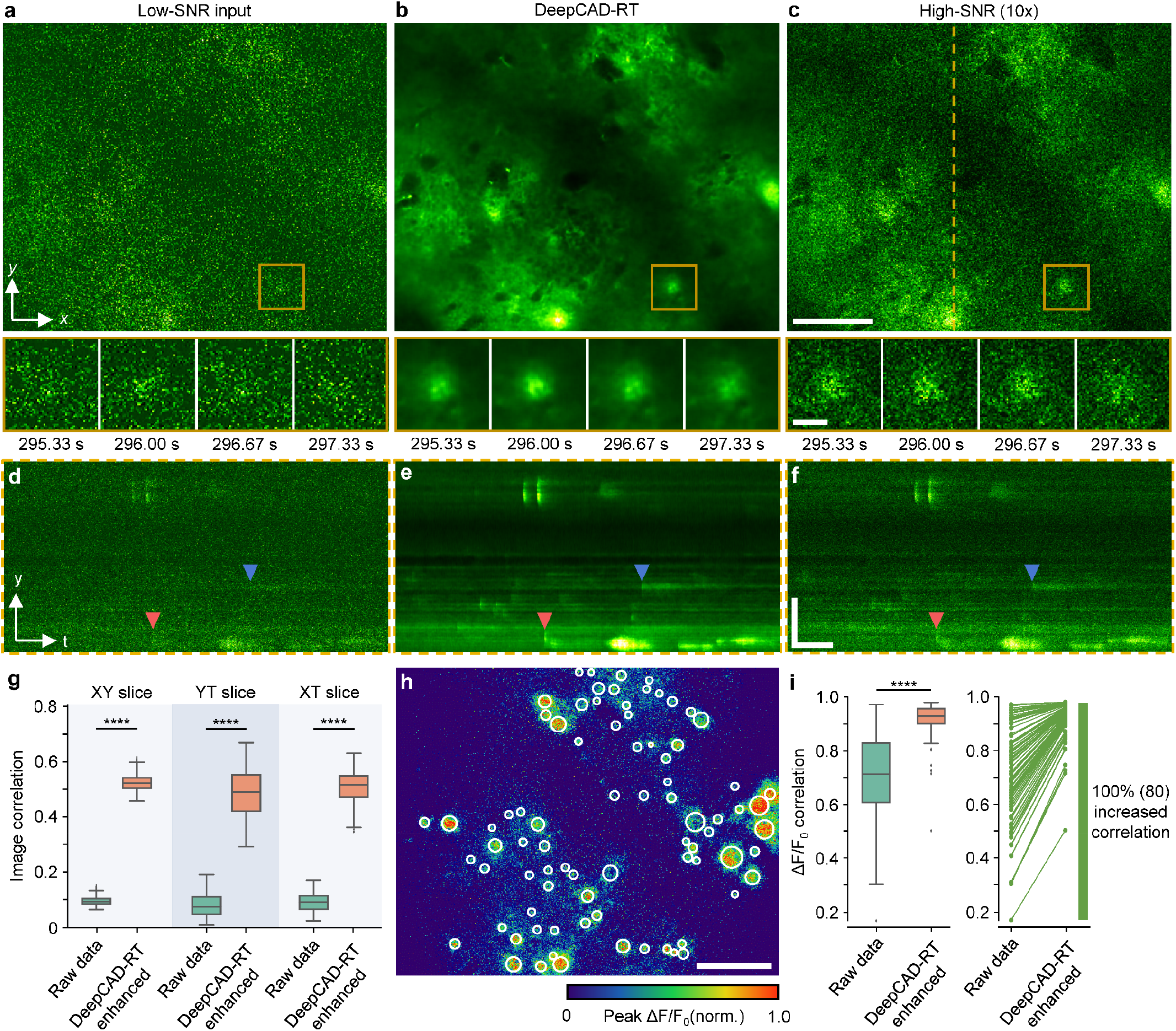
Denoising performance of DeepCAD-RT on neurotransmitter imaging in living mice. **a**, Low-SNR recording of extracellular ATP release in the mouse brain. **b**, DeepCAD-RT enhanced data with low-SNR recording as the input. **c**, Synchronized high-SNR data with 10-fold SNR. Magnified views showing ATP dynamics in the yellow boxed region in a 2-second period. Each frame was integrated for 67 ms. Scale bar, 50 μm for the large FOV and 10 μm for magnified views. **d-f**, YT slices along the dashed line in **c**. Two ATP-release events are indicated with arrowheads of different colors. Scale bar, 50 μm for y and 50 s for t. **g**, Pearson correlation coefficients of XY, YT, and XT slices before and after denoising. XY slice, N=7000; YT slice, N=476, XT slice, N=476. **h**, Peak ΔF/F_0_ of high-SNR data during the whole imaging session (~480 s). Manually annotated release sites are marked with white circles (N=80). Scale bar, 50 μm. **i**, Left, boxplots showing Pearson correlations of fluorescence traces extracted from release sites in **h** before and after denoising (N=80). High-SNR traces extracted from 10-fold SNR data were used as the ground truth for correlation calculation. Right, increases of trace correlation. Each line represents one of 80 traces and increased correlations are colored green. P values calculated by one-sided paired t-test are specified with asterisks. ****P < 0.0001 for all comparisons.

Previous studies about *in vivo* imaging of ATP release were restricted in 2D planes^33, 53^. To fully unveil the spatiotemporal distribution and evolution pattern of ATP release in 3D tissues, we performed volumetric imaging of a 350×350×60 μm^3^ tissue volume in the mouse brain after laser-ablated injury. The injury site was located at the center of the volume. Since inflammation and injury can trigger the release of endogenous ATP, phototoxicity and photodamage caused by the excitation laser should be minimized to avoid undesired disturbance. Thus, we kept the excitation power below 40 mW and imaged the 3D volume continuously for one hour. In the shot-noise-limited raw data, noise is dominant and only a few intense events can be seen (Fig. 5a). To suppress the shot noise and visualize as many release events as possible, we trained a denoising model with our method and then enhanced the original low-SNR data. Denoised data had very high SNR and those released events concealed by noise turned out to be discernable (Fig. 5a and Supplementary Video 10). For better comparison, we present several snapshots of a single plane at different moments (Fig. 5b,c), which indicates the superior denoising performance of our method. We manually annotated the position and time of all ATP-release events throughout the entire session (Fig. 5d) and found that the release frequency is approximately random during the one-hour imaging (Fig. 5e and Supplementary Fig. 13). Owing to the remarkable noise removal capability, the spatial profile of ATP release was clarified, and performing statistics on their geometric features (diameter and ellipticity) became feasible (Fig. 5f,g). The successful extension of DeepCAD-RT to the imaging of ATP release indicates its good potential on other neurotransmitter sensors.

**Fig. 5.**
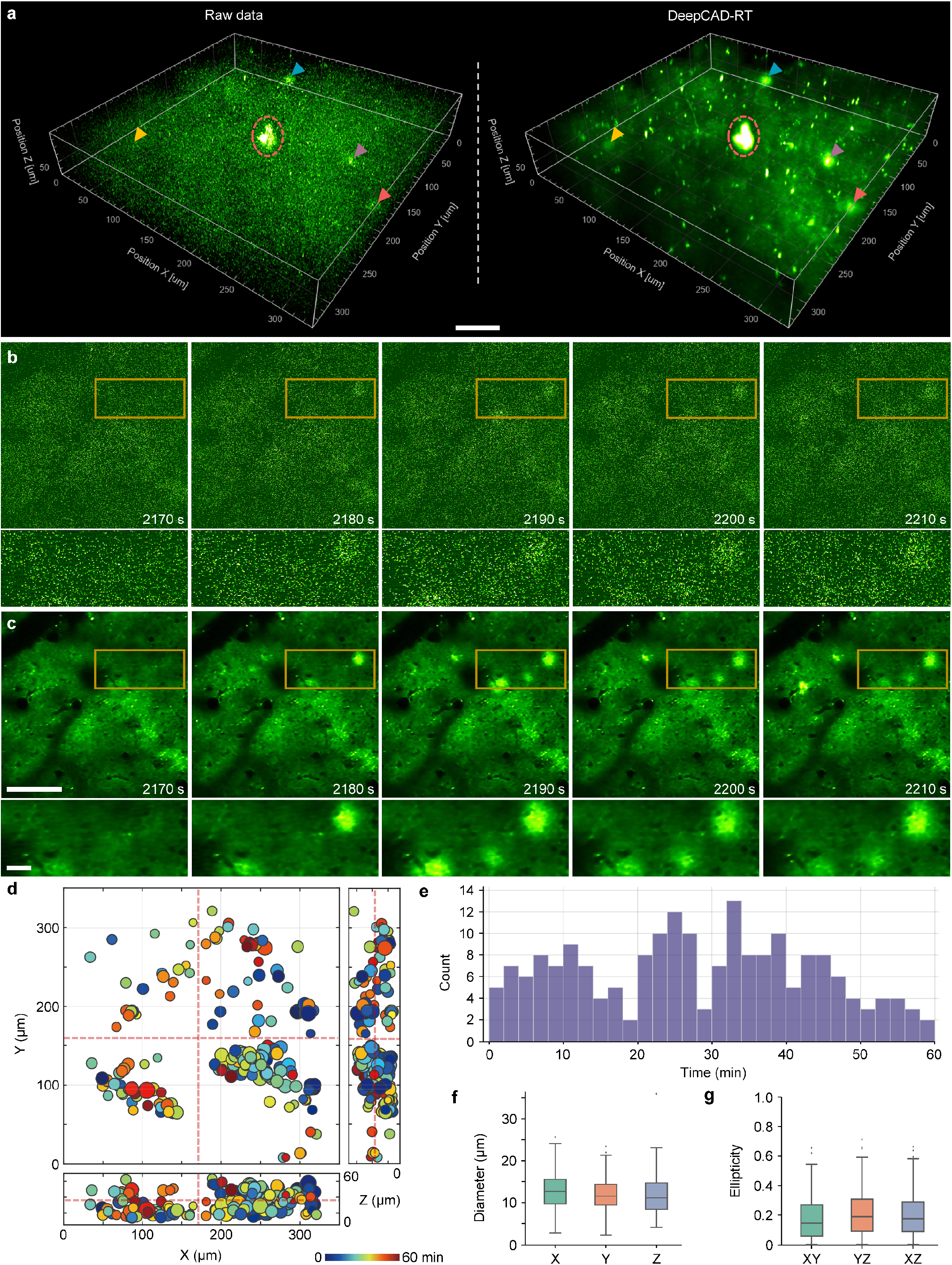
DeepCAD-RT reveals the spatiotemporal patterns of extracellular ATP *in vivo* after laser-induced brain injury. **a**, 3D visualization of ATP-release events in a 350×350×60 μm^3^ volume (30 planes, 1 Hz volume rate) after laser-induced brain injury. Left, low-SNR raw volume without denoising. Right, the same volume enhanced with DeepCAD-RT. A representative moment is demonstrated here and similar performance was achieved throughout the whole imaging session (1 hour, 3600 volumes). Four ATP-release events are indicated with arrowheads of different colors. The laser-ablated point (red dashed circle) was located at the center of the volume. Scale bar, 50 μm. **b**, Example raw frames of a single plane at four different time points. **c**, DeepCAD-RT enhanced frames corresponding to those in **b**. Magnified views of yellow boxed regions are shown under each image. Scale bar, 100 μm for the whole FOV and 20 μm for magnified views. **d**, The spatiotemporal distribution of ATP release during the one-hour-long recording. The release time is color-coded and the diameter of each release event scales to the size of each circle. The intersections of red dashed lines indicate the 3D location of the laser-induced injury. **e**, Counting ATP-release events along the time axis. The binning width is 2 min. **f**, Boxplots showing diameters of all release events (N=196) in three orthogonal dimensions. X, 13.131 ± 0.3090; Y, 12.125 ± 0.2911; Z, 11.907 ± 0.3287 (mean ± s.e.m.). **g**, Statistics on the ellipticity of all release events (N=196) in three orthogonal coordinate planes. XY, 0.182 ± 0.0109; YZ, 0.213 ± 0.0114; XZ, 0.205 ± 0.0109 (mean ± s.e.m.).

## Discussion

Noise is an ineluctable obstacle in scientific observation. For fluorescence microscopy, the inherent shot-noise limit determines the upper bound of imaging SNR and restricts imaging resolution, speed, and sensitivity. In this work, we present a versatile method to denoise fluorescence images with rapid processing speed that can be incorporated with the microscope acquisition system to achieve real-time denoising. Our method is based on deep self-supervised learning and the original low-SNR data can be directly used for training convolutional networks, making it particularly advantageous in functional imaging where the sample is undergoing fast dynamics and capturing ground-truth data is hard or impossible. We have demonstrated extensive experiments including calcium imaging in mice, zebrafish, and flies, cell migration observations, and the imaging of a new genetically encoded ATP sensor, covering both 2D single-plane imaging and 3D volumetric imaging. Qualitative and quantitative evaluations show that our method can substantially enhance fluorescence time-lapse imaging data and permit high-sensitivity imaging of biological dynamics beyond the shot-noise limit.

Removing shot noise from fluorescence images promises to catalyze advancements in several imaging technologies. For example, in two-photon microscopy, multiplexed excitation by multiple laser foci can increase imaging speed but the imaging SNR will decrease quadratically because of dispersed excitation power^54–56^. Our denoising method provides a potential solution to compensate for the SNR loss. Three-photon microscopy can effectively suppress background fluorescence and improve imaging depth through three-order nonlinear excitation and longer wavelength^57, 58^, but its practical use in deep tissue is still limited by low imaging SNR. Combining our method with three-photon microscopy could expedite its application in the deep mammalian brain. Light-field microscopy is an emerging technique for fast volumetric imaging of biological dynamics, but it relies on computational reconstruction that is sensitive to noise^59–61^. Disentangling underlying signals from noisy images before light-field reconstruction could eliminate artifacts and ensure high-fidelity results. Moreover, a recently published work reported that standard Richardson–Lucy deconvolution can recover high-frequency information beyond the spatial frequency limit of the microscope if there is no noise contamination^62^, which inspires us that our method would be helpful for deconvolution algorithms by denoising input images in advance. Single-molecule localization microscopy (SMLM) is also susceptible to noise since the localization precision is fundamentally limited by SNR^3, 63^. The noise-sensitive nature holds for other super-resolution microscopy techniques such as stimulated emission depletion (STED) microscopy and structured illumination microscopy (SIM)^64, 65^. We reasonably envisage that our method and its future variants would benefit the development of super-resolution microscopy.

As the core of our method lies in deep learning, its content-dependent trait requires users to train a specialized model for each task or each type of sample to ensure optimal results. Developing pre-trained models on large-scale datasets and then transferring them to new tasks by fine-tuning could be an optional solution to this problem. Another limitation is that adjacent frames used for training should have approximately identical underlying signals, which is the basic assumption of our self-supervised training strategy. Thus, the imaging system should have adequate temporal resolution relative to the biological dynamics to be imaged. Finally, the denoising performance of our method improves as the SNR of the input data increases. Comprehensive noise suppression by collaborating physics-based approaches^19, 28^ and computational denoising could be a more powerful way to break the shot-noise limit.

## Methods

### Imaging system

The optical setup integrated two two-photon microscopes for different purposes. One was a standard two-photon microscope with multi-color detection capability for multi-labeling imaging and cross-system validation. The other one was a custom-designed two-photon microscope to capture synchronized low-SNR and high-SNR (10-fold SNR) images for result validation (Supplementary Fig. 9). The two systems shared one femtosecond titanium-sapphire laser source with tunable wavelength (Mai Tai HP, Spectra-Physics). The excitation laser for all experiments was a linearly polarized Gaussian beam with a 920-nm central wavelength and an 80-MHz repetition rate. Before being projected into both systems, the laser beam was first adjusted in polarization by a half-wave plate (AQWP10M-980, Thorlabs) and modulated in intensity by an electro-optic modulator (350-80LA-02, Conoptics). A 1:1 4f system composed of two achromatic convex lenses (AC508-100-B, Thorlabs) was then configured to collimate the laser beam. Another 1:4 4f system (AC508-100-B and AC508-400-B, Thorlabs) was followed to expand the diameter of the beam. A mirror mounted on a two-position, motorized flip mount (MFF101, Thorlabs) was used to alternate between the two systems (OFF for the multi-color module and ON for the custom module).

The two systems used the same optical configuration for two-photon excitation. Specifically, the collimated, scaled laser beam was successively guided onto the fast axis (the resonant mirror) and the slow axis (the galvanometric mirror) of the galvo-resonant scanner (8315K/CRS8K, Cambridge Technology). The scanner provided fast 2D raster scanning under the control of two voltage signals. The orientation of the incident beam should be fine-adjusted to ensure the horizontality of the outgoing beam. Then, the output beam was recollimated, rescaled, and corrected by a scan lens (SL50-2P2, Thorlabs) and a tube lens (TTL200MP, Thorlabs) to fit the back pupil of the objective and produce a flat image plane. We used a high-numerical-aperture (NA) water-immerse objective (×25/1.05 NA, XLPLN25XWMP2, Olympus) to expand the detection angle and increase the number of photons that can be detected. Approximately, the effective excitation NA was 0.7 in our experiments. To perform 3D volumetric imaging, we mounted the objective on a piezoelectric actuator (P-725, Physik Instrumente) to achieve high-precision axial scanning. For the detection path of the standard multi-color system, fluorescence photons emitted from the sample were captured by the objective and then separated from the excitation light by a long-pass dichroic mirror (DMLP650L, Thorlabs). Another short-pass dichroic mirror (DMSP550, Thorlabs) was mounted in the detection path to separate green fluorescence and red fluorescence. The green fluorescence was purified by a pair of emission filter (MF525-39, Thorlabs; ET510/80M, Chroma) and then detected by a GaAsP photomultiplier tube (H10770PA-40, Hamamatsu). The red fluorescence was filtered by an emission filter (ET585/65M, Chroma) and then detected by the same type of PMT. For the detection path of the customized system for simultaneous low-SNR and high-SNR imaging, the previously mentioned short-pass dichroic mirror was replaced with a 1:9 (reflectance: transmission) non-polarizing plate beam splitter (BSN10, Thorlabs). Low-SNR images were formed by the ~10% reflected photons and high-SNR images were formed by the ~90% transmitted photons. In this system, only green fluorescence was detected and the same filters and PMT were used for both the low-SNR and high-SNR detection path. The sensor plane of each PMT was conjugated to the back-pupil plane of the objective using a 4:1 4f system (TTL200-A and AC254-050-A, Thorlabs) to maximize the detection efficiency. In general, the maximum FOV of the two two-photon microscopes was about 720 μm. The typical frame rate was 30 Hz for 512×512 pixels and the volume rate decreased linearly with the number of planes to be scanned.

### System calibration

We imaged green-fluorescent beads to calibrate our imaging systems. For sample preparation, the original bead suspension was first diluted and embedded in 1.0% agarose and then mounted on microscope slides to form a single bead layer composed of sparsely distributed beads. We calibrated both systems using 0.2-μm fluorescent beads (G200, Thermo Fisher) to obtain the lateral and axial resolution. Since the two systems had identical excitation optics, they had the same optical resolution. The lateral full width at half maximum (FWHM) is ~0.6 μm and the axial FWHM is ~3.5 μm (Supplementary Fig. 14). To calibrate the intensity ratio between the high-SNR detection path and the low-SNR detection path, we imaged 1-μm fluorescent beads (G0100, Thermo Fisher) and found that the intensity ratio is about 1:10 (Supplementary Fig. 10a-d), which indicated that the imaging SNR of the high-SNR detection path was about ten times higher than that of the low-SNR detection path. High-SNR data synchronized with low-SNR data could serve as a reference to unveil underlying signals. We also imaged insect slices for validation and the results confirmed our calibration (Supplementary Fig. 10e-h).

### Model simplification

Theoretically, large models with more trainable parameters can implement extremely intricate functions on the input data. However, the very big model (16,315,585 (16.3 M for short) parameters in total) we previously used caused a series of problems such as long training and inference time, large memory consumption, and serious overfitting. We sought to solve these problems by simplifying the network architecture. Since network depth is of crucial importance for the performance^66^, instead of changing the depth of the network, we turned to reduce the number of feature maps in each convolutional layer. By continuously halving network parameters, we constructed five models with exponentially decreased trainable parameters (16.3 M, 9.2 M, 4.1 M, 2.3 M, 1.0 M, respectively). To evaluate these models, we used synthetic calcium imaging data of −2.5 dB SNR and trained them with the same amount of data (6000 frames). The best training epoch of each model was determined by monitoring its performance on a holdout validation set. Although the number of trainable parameters was reduced by ~94%, the denoising performance remained almost unchanged (Supplementary Fig. 2). A more comprehensive assessment including training and inference time, memory consumption, and output SNR is shown in Supplementary Table 2. The lightweight model with ~1.0-million parameters was chosen as the final architecture.

### Data augmentation

The strategy to eliminate overfitting by drastically reducing trainable parameters only works when there is enough training data. If only a small dataset is available, overfitting still occurs even with very small models^67^. To alleviate the data dependency of our method and further eliminate overfitting, we designed 12-fold data augmentation to generate enough training pairs from a small amount of data (Supplementary Fig. 4). Given a low-SNR time-lapse image stack, thousands of 3D training pairs with overlaps will be extracted from the input stack. A training pair includes an input patch and a corresponding target patch. The proportion of temporal overlapping was automatically calculated according to the number of training pairs to be extracted. For each training pair, we first swapped the input and target randomly with a probability of 0.5. Then, we performed six geometric transformations randomly for the training pair including horizontal flip, vertical flip, left 90-degree rotation, 180-degree rotation, right 90-degree rotation, and no transformation. Overall, there were 12 possible forms for each training pair and they all have the same probability of occurrence, which inflated the training dataset by 12-fold. We investigated the benefit of our data augmentation strategy using synthetic calcium imaging data and found that the data dependency of our method was reduced effectively (Supplementary Fig. 5). A 1000-frame calcium imaging stack (490×490 pixels) is enough to train a model with satisfactory performance. This feature is helpful to alleviate the problem of insufficient training data in fluorescence microscopy. To evaluate the effect of data augmentation on overfitting, we trained a model with data augmentation and the other one without data augmentation with the same amount of data for a long training period (35 epochs) and monitored their performance after each epoch. The results show that training with data augmentation could keep the performance stable compared to the rapidly degrading performance without augmentation (Supplementary Fig. 6). The optimal performance was also improved because of augmented training data. Although the combination of model simplification and data augmentation eliminates overfitting, preparing more training data is still the most effective way to improve the denoising performance and avoid overfitting.

### Network architecture, training and inference

The network architecture in this research reserves the topology of 3D U-Net^68^ that utilizes the encoder-decoder paradigm in an end-to-end manner. To fully exploit spatiotemporal correlations in fluorescence imaging data, all operations inside the network were implemented in 3D, including convolutions, max-poolings, and interpolations (Supplementary Figure 14). Compared to our previous architecture^32^, the number of feature maps in each convolutional layer was reduced by 4-fold and the total number of trainable parameters was reduced by 16-fold (1,020,337 compared with 16,315,585), which massively improved the training and inference speed and reduced the memory consumption. For pre-processing, each input stack was subtracted by the average of the whole stack to handle the intensity variation across different samples and imaging platforms. These stacks were partitioned into a specified number of 3D (x-y-t) training pairs. The data augmentation strategy mentioned above would be applied to each training pair. Training was carried out using the arithmetic average of an L1-norm loss term and an L2-norm loss term as the loss function. After the input stack flowed through the network, the subtracted average value would be added back in post-processing. Since the combination of model simplification and data augmentation eliminated overfitting, the model of the last training epoch could be directly selected as the final solution. For denoising of 3D volumetric imaging, the time-lapse stack of each imaging plane was saved as a separate TIFF file. All stacks were used for the training of the network.

The batch size for all experiments was set to the number of GPUs being used. The patch size was set to 150×150×150 pixels by default. All models were trained using the Adam optimizer^69^ with a learning rate of 5×10^-5^, and the exponential decay rates for the first-moment and second-moment estimates were 0.5 and 0.9, respectively. Using our Python code, training with 3000 pairs of 3D patches for 20 epochs just took 6.2 hours on a single GPU (GeForce RTX 3090, Nvidia). The inference process for an image stack composed of 490×490×300 pixels (partitioned into 75 3D patches) took as few as 8 seconds. Multi-GPU acceleration has been supported by our Python code. The time consumption of training and inference decreases linearly as the number of GPUs increases.

### Real-time implementation of DeepCAD-RT

To achieve real-time processing during imaging acquisition, we made a program interface to incorporate DeepCAD-RT into our image acquisition software (Scanimage 5.7^70^, Vidrio Technologies). For further acceleration and memory conservation, the inference of DeepCAD-RT was optimally deployed on GPU with TensorRT (NVIDIA), a software development kit providing low-latency and high-throughput processing for deep learning applications by executing customized operation automatically for specific GPU and network architecture. Three parallel threads were designed for imaging, data processing, and display. The schedule for multi-thread programming is depicted in Fig. 1c. Specifically, the first thread was used for image acquisition, which waited for a certain number of frames and packaged them into 3D (x-y-t) batches. Adjacent batches had overlapping frames and half of the overlap would be discarded to avoid artifacts. Then, the second thread got low-SNR images passed by the first thread, processed them, and produced denoised frames. Finally, these denoised frames were transferred to the third thread for display. When the imaging process stopped, denoised images would be automatically saved in a user-defined directory. The real-time implementation was programmed in C++ for best hardware interaction and then compiled in Matlab (MathWorks), which could be called by any Matlab-based software or script. On a single GPU (GeForce RTX 3090, Nvidia), the real-time implementation achieved more than 20-fold speed-up compared to the original DeepCAD^32^ and had an extremely low memory consumption as few as 701 MB with float16 precision. The real-time implementation of DeepCAD-RT has been packaged as a free plugin with a user-friendly interface (Supplementary Fig. 3). To transfer pre-trained models, scripts was developed to convert PyTorch models to ONNX (Open Neural Network Exchange) models and then call TensorRT builder to optimize ONNX models for a target GPU, which produced engine files that can be used by TensorRT. The construction of the engine file would eliminate dead computations, fold constants, and combine operations to find an optimal schedule for model execution.

### Animal preparation and fluorescence imaging

Multiple animal models (mouse, zebrafish, and fly) and fluorescence labeling methods (calcium, neutrophils, ATP release) were associated in this research. All experiments involving animals were performed in accordance with the institutional guidelines for animal welfare and have been approved by the Animal Care and Use Committee of Tsinghua University.

#### Mouse preparation and imaging

Adult mice (male or female without randomization or blinding) at 8–16 postnatal weeks were housed in animal facility (24 °C, 50% humidity) under a reverse light cycle in groups of 1–5. All imaging experiments were carried out with our two-photon microscopes on head-fixed, awake mice.

For functional imaging of neural activity, we used transgenic mice hybridized between Rasgrf2-2A-dCre mice and Ai148 (TIT2L-GC6f-ICL-tTA2)-D mice expressing Cre-dependent GCaMP6f genetically encoded calcium indicator (GECI). Craniotomy surgeries were conducted for chronic two-photon imaging as previously described^32^. Briefly, mice were first anesthetized with 1.5% (by volume in O_2_) isoflurane and a 6.0-mm diameter craniotomy was made with a skull drill. After removing the skull piece, a coverslip was implanted on the craniotomy region and a titanium headpost was then cemented to the skull for head fixation. After the surgery, 0.25 mg/g (body weight) trimethoprim (TMP) was injected intraperitoneally to induce the expression of GCaMP6f in layer 2/3 cortical neurons across the whole brain. After the inflammation was gone and the cranial window became clear (~2 weeks after surgery), mice were head-fixed on a customized holder with a 3D-printed plastic tube to restrict the mouse body. The holder was mounted on a high-precision, three-axis motorized stage (M-VP-25XA-XYZL, Newport) for sample translation. *In vivo* calcium imaging (30-Hz single-plane imaging) was carried out on awake mice without anesthesia. The imaging of dendritic spines in cortical layer 1 (20-60 μm below the brain surface) required adequate spatial sampling rate that was achieved by using large zoom factors.

For time-lapse imaging of neutrophil migration, we first performed craniotomy on wild-type mice (C57BL/6J) following the procedures described above. Acute brain injury caused by craniotomy would induce immune responses in the brain. After the surgery, neutrophils and blood vessels were simultaneously labeled by injecting 10 μg red (Alexa Fluor 555 conjugate) wheat germ agglutinin (WGA) dye (W32464, Thermo Fisher Scientific) and 2 μg of green-fluorescence-conjugated Ly-6G/Ly-6C antibody (53-5931-82, eBioscience) intravenously. The two dyes were dissolved and diluted in 200 μL 1× phosphate-buffered saline (PBS). To avoid the potential influence of anesthesia on immune response, *in vivo* two-photon imaging was performed in the mouse brain after the mouse was fully awake (~20 minutes after injection). Imaging experiments should be finished as soon as possible since these dyes are degradable in the mouse body. Empirically, the whole imaging session should take no longer than 5 hours. Volumetric imaging was implemented by scanning the objective axially with the piezoelectric actuator. The frame rate of single-plane imaging was 30 Hz and the volume rate of 3D imaging was 2 Hz (15 imaging planes). The whole 3D imaging session lasted ~20 minutes. For each 3D volume, the flyback frame acquired while the piezoelectric actuator was quickly returning from the bottom plane to the top plane should be discarded. Images of the green channel and the red channel were captured simultaneously and were separated by post-processing.

For functional imaging of ATP dynamics, wild-type mice (C57BL/6J) were anesthetized with intraperitoneally injected Avertin (500 mg/kg body weight, Sigma-Aldrich). A cranial window was opened on the visual cortex and 400-500 nL AAV (AAV2/9-GfaABC1D-ATP1.0, packaged at Vigene Biosciences) was injected (AP: −2.2 mm relative to Bregma, ML: 2.0 mm relative to Bregma, and DV: 0.5 mm below the dura, at an angle of 30°) using a micro-syringe pump (Nanoliter 2000 injector, World Precision Instruments) to express GRAB_ATP1.0_^33^ in cortical astrocytes. A 4 mm × 4 mm square coverslip was implanted to replace the skull. After ~3 weeks of recovery and virus expression, two-photon imaging was performed to record ATP-release events in the mouse cortex. Before imaging, brain injury was induced by ablating the tissue with a stationary laser focus (200 mW) for 5 seconds. The injury site was located at the center of the 3D imaging volume. Single-plane images were recorded at the plane 20 μm above the injury site. The frame rate of single-plane imaging was 30 Hz and the volume rate of 3D imaging was 1 Hz (30 imaging planes). The flyback frame of each volume should be discarded. Only signals from the green channel were recorded and the whole 3D imaging session lasted 60 minutes.

#### Zebrafish preparation and imaging

Transgenic zebrafish (*Danio rerio*) larvae expressing pan-neuronal GCaMP6s calcium indicator (Tg(HuC:GCaMP6s)) were housed in culture dishes at 28.5 °C in Holtfreter’s solution (59 mM NaCl, 0.67 mM KCl, 0.76 mM CaCl_2_, 2.4 mM NaHCO_3_). At 4-6 days postfertilization (dpf), zebrafish larvae were separated and restricted in a small drop of 1.0% low melting point agarose (Sigma-Aldrich) and then mounted on a microscope slide for imaging. A fine-bristle brush was used to adjust the posture of the larvae to keep the dorsal side up before the agarose solidified. After fixation, the larvae were placed under the objective and Holtfreter’s solution was used as the immersion medium of the objective. Before image acquisition started, we previewed the image and rotated the microscope slide manually to keep the larva horizontal or vertical in the FOV. Two-photon calcium imaging of spontaneous neural activity was performed on the larvae at 26–27 °C without anesthesia or motion paralysis. All experiments were single-plane imaging and the frame rate was 30 Hz for 512×512 pixels. Both large neuronal populations across multiple brain regions and small neuronal subsets localized in the optic tectum were imaged using different zoom factors.

#### Drosophila preparation and imaging

Flies were raised on standard cornmeal medium with a 12h/12h light/dark cycle at 25°C. Transgenic flies UAS-GCaMP7f were crossed with OK107-Gal4 to drive the expression of GCaMP7f^24^ calcium indicator in essentially all Kenyon Cells.

All experiments were conducted on female F1 heterozygotes from this cross. Flies at 5 days posteclosion were anesthetized on ice and mounted in a 3D-printed plastic disk that allowed free movement of the legs as previously reported^71^. The posterior head capsule was opened using sharp forceps (5SF, Dumont) under room temperature in carbonated (95% O_2_, 5% CO_2_) buffer solution (103 mM NaCl, 3 mM KCl, 5mM N-Tris, 10 mM trehalose, 10 mM glucose, 7mM sucrose, 26 mM NaHCO_3_, 1mM NaH_2_PO_4_, 1.5 mM CaCl_2_, 4mM MgCl_2_) with a pH of 7.3 and an osmolarity of 275 mOsm. After that, the air sacks and tracheas were also removed. Brain movement was minimized by adding UV glue around the proboscis and removing the M16 muscle^37, 72^. After the preparation, flies were placed under the objective for two-photon imaging of calcium transients in the mushroom body. To enhance the neural activity, 4-methylcyclohexanol (MCH) and 3-octanol (OCT) 1:1000 diluted in mineral oil (MO) were used as odors. Flies were randomly given the two odors for five seconds every ten seconds using a custom-made air pump. All experiments were single-plane imaging at 30 Hz with 512×512 pixels.

### Generation of synthetic calcium imaging data

We used synthetic calcium imaging data (simulated time-lapse image sequences) for quantitative evaluations of our method, as well as for comparisons with DeepInterpolation^31^. Our simulation pipeline consisted of synthesizing noise-free calcium imaging videos (ground truth) and adding different levels of Mixed Poisson-Gaussian (MPG) noise^21, 32^ to them. To generate noise-free calcium imaging data, we adopted in silico Neural Anatomy and Optical Microscopy (NAOMi), a simulation method to create realistic calcium imaging datasets for assessing two-photon microscopy methods^35^. The parameters of our simulation are listed in Supplementary Table 2. Those not mentioned all used default values. Simulated data had very similar spatiotemporal features as experimentally obtained data including neuronal anatomy (cell bodies, neuropils, dendrites, etc.), neural activity, and blood vessels. For noise simulation, we first performed Poisson sampling on noise-free images to simulate the content-dependent Poisson noise. Then we added content-independent Gaussian noise to these data. Poisson noise was set as the dominant noise source. Different imaging SNRs were simulated by different relative photon numbers that changed the intensity of input noise-free images (Supplementary Fig. 1).

### Neutrophil segmentation

Four types of data were involved in this experiment, *i.e*., raw data (low-SNR), high-SNR (10× SNR) data, denoised raw data, and denoised high-SNR data. Ten representative images with relatively sparse cells were selected from the dataset of single-plane neutrophil imaging for semantic segmentation. To obtain ground-truth segmentation masks, five human experts were recruited to annotate all neutrophils in each denoised high-SNR image using the ROI Manager toolbox of Fiji. The final ground-truth masks were determined by majority voting. Neutrophil segmentation was conducted using Cellpose^43^ and Stardist^44^, two CNN-based, generalist algorithms for cellular segmentation. For both methods, default parameters and pre-trained models were used without additional training. Segmentation performance was quantitatively evaluated with the Intersection-over-Union (IoU) score^73^ defined as

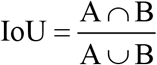

where A is the mask segmented by algorithms and B is the ground truth. Statistical analysis and representative results were summarized in Supplementary Fig. 12.

### 3D visualization

For volumetric imaging of neutrophil migration and ATP release, we performed 3D visualization to reveal the spatiotemporal patterns of biological dynamics. Imaris 9.0 (Oxford Instruments) was used for the visualization of all volumetric imaging data. Both the original low-SNR data and denoised data were imported into Imaris, rendered with pseudo-color, and 3D reconstructed using the maximum intensity projection mode. The brightness of data before and after denoising was adjusted to make them have a similar visual effect. The contrast of low-SNR data was fine-tuned to show underlying signals as clearly as possible. All values for gamma correction were set to one. The red channel (blood vessels) of neutrophil migration was averaged by multiple frames to improve its SNR and then merged with the green channel. Crosstalk signals out of the blood vessel were manually suppressed with Fiji. Animations were generated by automatically interpolating intermediate frames between selected keyframes.

### Annotation of ATP-release events

The whole annotation pipeline was implemented on the denoised data (Supplementary Figure 13). The spatial shape of each ATP-release event could be modeled as an ellipsoid. To obtain the center position and peak time of each event throughout the whole imaging session, we manually annotated them by adding measurement points in Imaris. All spatial and temporal coordinates were exported from the software after annotation. Events at the edge of the volume were excluded because only a part of them appeared in the FOV. Based on these annotated coordinates, intensity profiles along all three dimensions of each event were extracted from denoised stacks with a custom Matlab (MathWorks) script. Gaussian fitting was performed for all intensity profiles to reduce the influence of background fluctuations. Then, all fitted Gaussian curves were deconvolved with the system point spread function (PSF) (Supplementary Figure 15) using standard Richardson–Lucy algorithm^74, 75^. This step eliminated the influence of limited and anisotropic spatial resolution. The diameter of these ATP-release events could be extracted in each dimension, which was defined as the FWHM of deconvolved gaussian curves. The ellipticity of release events was defined as

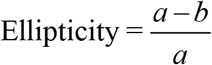

where *a* is the major axis of the ellipse and *b* is the minor axis of the ellipse. Ellipticity was calculated for each 3D release event in all three orthogonal coordinate planes (XY, YZ, XZ).

### Performance metrics

To quantitatively evaluate the performance of our method, both synthetic data and experimentally obtained data were used. For synthetic calcium imaging data, ground-truth images were available and SNR was calculated to quantify the denoising performance. SNR was defined as the logarithmic form:

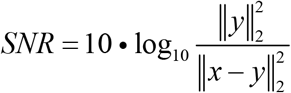

where *x* is the denoised data and *y* is the ground truth. For experimentally obtained data, synchronized high-SNR data with 10-fold SNR acquired with our system were used as the reference of underlying signals. Pearson correlation coefficient (R) was used as the performance metric, which is formulated as

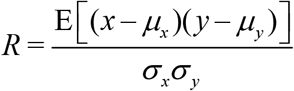

where *x* and *y* are the denoised data and corresponding high-SNR data, respectively; *μ_x_* and *μ_y_* are the mean values of *x* and *y*; *σ_x_* and *σ_y_* are the standard deviations. The operator E represents arithmetically averaging. Pearson correlation was used for both images and fluorescence traces. All performance metrics were implemented with custom Matlab scripts and built-in functions.

### Statistics and reproducibility

Sample sizes and statistics are reported in the figure legends and text for each experiment. All boxplots were plotted in the format of standard Tukey box-and-whisker plot. The box indicates the lower and upper quartiles while the line in the box shows the median. The lower whisker represents the first data point greater than the lower quartile minus 1.5× the interquartile range (IQR). Similarly, the upper whisker represents the last data point less than the upper quartile plus 1.5× the IQR. Outliers were plotted in small black dots. For the comparison of images and fluorescence traces before and after denoising, one-sided paired t-test was performed and P values were indicated with asterisks. Representative frames were demonstrated in the figures and similar results were achieved on more than 1500 frames for all experiments.

## Data availability

We have no restriction on data availability. All source data (~250 GB), including synthetic calcium imaging data, experimental recordings of calcium dynamics, neutrophil migration, and cortical ATP release, have been archived and made publicly available at https://cabooster.github.io/DeepCAD-RT/Datasets/.

## Code availability

All relevant resources are readily accessible on our GitHub page https://cabooster.github.io/DeepCAD-RT/. The source PyTorch code, demo notebooks (in Jupyter Notebook and Google Colab), and the code for real-time implementation can be found at https://github.com/cabooster/DeepCAD-RT/. A detailed tutorial for all codes has been provided at https://cabooster.github.io/DeepCAD-RT/Tutorial/.

## Acknowledgements

We would like to acknowledge Z. Jiang for providing the zebrafish larvae used in this research, and D. Jiang for providing dyes used for neutrophil labeling. We thank Z. Wang and R. Zhang for their support in mouse surgery. This work was supported by the National Natural Science Foundation of China (62088102, 62071272, 61831014, 62125106), the National Key Research and Development Program of China (2020AA0105500), and the Shenzhen Science and Technology Project under Grant (CJGJZD20200617102601004, ZDYBH201900000002). We further thank the supports from Beijing Laboratory of Brain and Cognitive Intelligence, Beijing Municipal Education Commission, and Beijing Key Laboratory of Multi-dimension & Multi-scale Computational Photography (MMCP).

## Author Contributions

Q. D., H. W. and L. F. supervised this research. Q. D., H. W., L. F. and X. L. conceived and initiated this project. X. L. designed detailed implementations, built the imaging system, and performed imaging experiments under the instruction of J. W., H. W., L. F. and Q. D. X. L. and YX. L. developed the Python code, performed simulations, and processed relevant imaging data. YX. L., Y. Zhou. and X. L. developed the real-time implementation. J. W., Y. Zhou, Z. Z., J. F., G. X., J. H., Y. Zhang, G. Z., H. X, and H. Q. gave critical support on system setup and imaging procedure. J. F., G. X, J. H., F. D., Z. W. and Y. L. provided animal models and prepared samples. X. L., YX. L, Y. Zhou, Z. Z. and X. H. annotated masks of neutrophil segmentation. X. L. and YX. L. analyzed the data, prepared figures and videos, and made the companion webpage. X. L., J. W., Y. Zhang, F. D., Z. W., X. H., Y. L., H. W., L. F. and Q. D. participated in discussions about the results. All authors participated in the drafting of the manuscript.

## Materials & Correspondence

Correspondence and requests for materials should be addressed to H. W., L. F. or Q. D.

