## Supplementary Information for "Real-time denoising of fluorescence time-lapse imaging enables high-sensitivity observations of biological dynamics beyond the shot-noise limit"

### CONTENT

#### I. Supplementary Figures


| | |
|---|---|
| **Supplementary Figure 1** | *In silico* simulation of calcium imaging data. |
| **Supplementary Figure 2** | Evaluating the performance of different model complexity. |
| **Supplementary Figure 3** | Real-time implementation of DeepCAD-RT. |
| **Supplementary Figure 4** | Data augmentation strategy. |
| **Supplementary Figure 5** | Evaluating the data dependency of DeepCAD-RT. |
| **Supplementary Figure 6** | Training stability with and without data augmentation. |
| **Supplementary Figure 7** | Comparing DeepCAD-RT and DeepInterpolation at different noise levels. |
| **Supplementary Figure 8** | Performance comparison between DeepCAD-RT and DeepInterpolation on neutrophil imaging data. |
| **Supplementary Figure 9** | Imaging system. |
| **Supplementary Figure 10** | System calibration. |
| **Supplementary Figure 11** | Denoising calcium imaging across multiple brain regions in zebrafish. |
| **Supplementary Figure 12** | The performance of neutrophil segmentation before and after denoising. |
| **Supplementary Figure 13** | ATP annotation pipeline. |
| **Supplementary Figure 14** | Network architecture. |
| **Supplementary Figure 15** | System point spread function (PSF). |


#### II. Supplementary Tables


| | |
|---|---|
| **Supplementary Table 1** | Comparison of different model complexity. |
| **Supplementary Table 2** | Parameters for the simulation of calcium imaging data |

#### III. Supplementary Videos

| | |
|---|---|
| **Supplementary Video 1** | Demonstrating real-time denoising on a two-photon microscope using DeepCAD-RT. |
| **Supplementary Video 2** | DeepCAD-RT enhances the *in vivo* recording of calcium transients in dendritic spines. |
| **Supplementary Video 3** | DeepCAD-RT massively improves the imaging SNR of neuronal population recordings in the zebrafish brain. |
| **Supplementary Video 4** | DeepCAD-RT massively improves the imaging SNR of neuronal population recordings across multiple brain regions in the zebrafish brain. |
| **Supplementary Video 5** | DeepCAD-RT enhances the neuronal population imaging of Drosophila mushroom body. |
| **Supplementary Video 6** | Denoising performance of DeepCAD-RT on two-photon imaging of neutrophils in the mouse brain. |
| **Supplementary Video 7** | DeepCAD-RT facilitates high-SNR observations of retraction fiber dynamics during neutrophil migration. |
| **Supplementary Video 8** | DeepCAD-RT reveals the 3D migration of neutrophils *in vivo* after acute brain injury. |
| **Supplementary Video 9** | Denoising performance of DeepCAD-RT on a recently developed genetically encoded ATP (Adenosine 5'-triphosphate) sensor. |
| **Supplementary Video 10** | DeepCAD-RT reveals the ATP (Adenosine 5'-triphosphate) dynamics of astrocytes in 3D after laser-induced brain injury. |

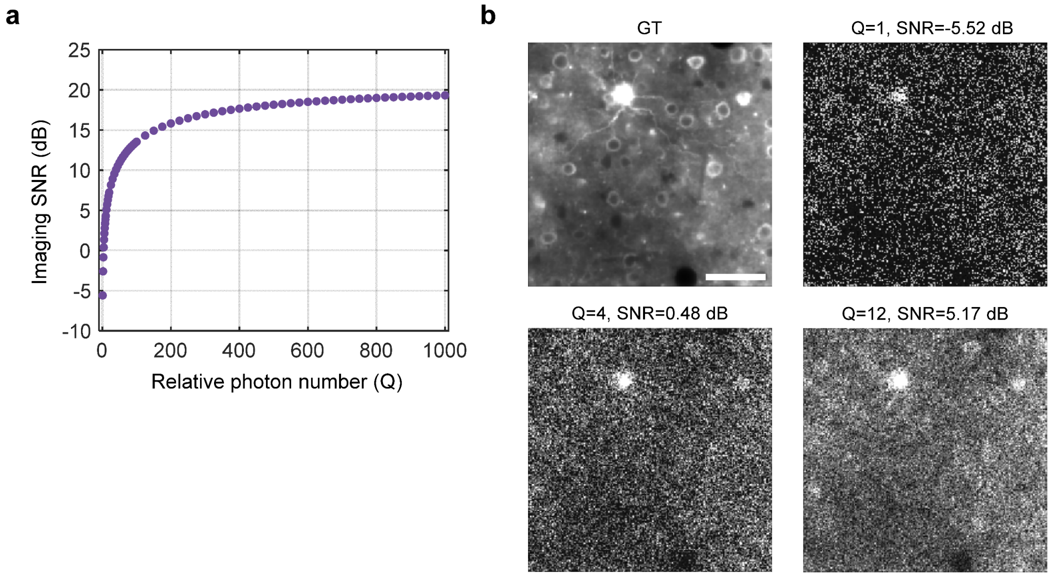


**Supplementary Figure 1**

**In silico simulation of calcium imaging data.**

Noise-free two-photon calcium imaging videos were simulated with *in silico* Neural Anatomy and Optical Microscopy (NAOMi)[1]. Different levels of Mixed Poisson-Gaussian (MPG) noise[2, 3] were added subsequently. Noise-free images were used as the ground truth for quantitative evaluations of the denoising performance. **a,** The relation between imaging SNR and relative photon number (Q). **b,** Representative images of different imaging SNRs and corresponding ground truth (GT). Scale bar, 50 μm.

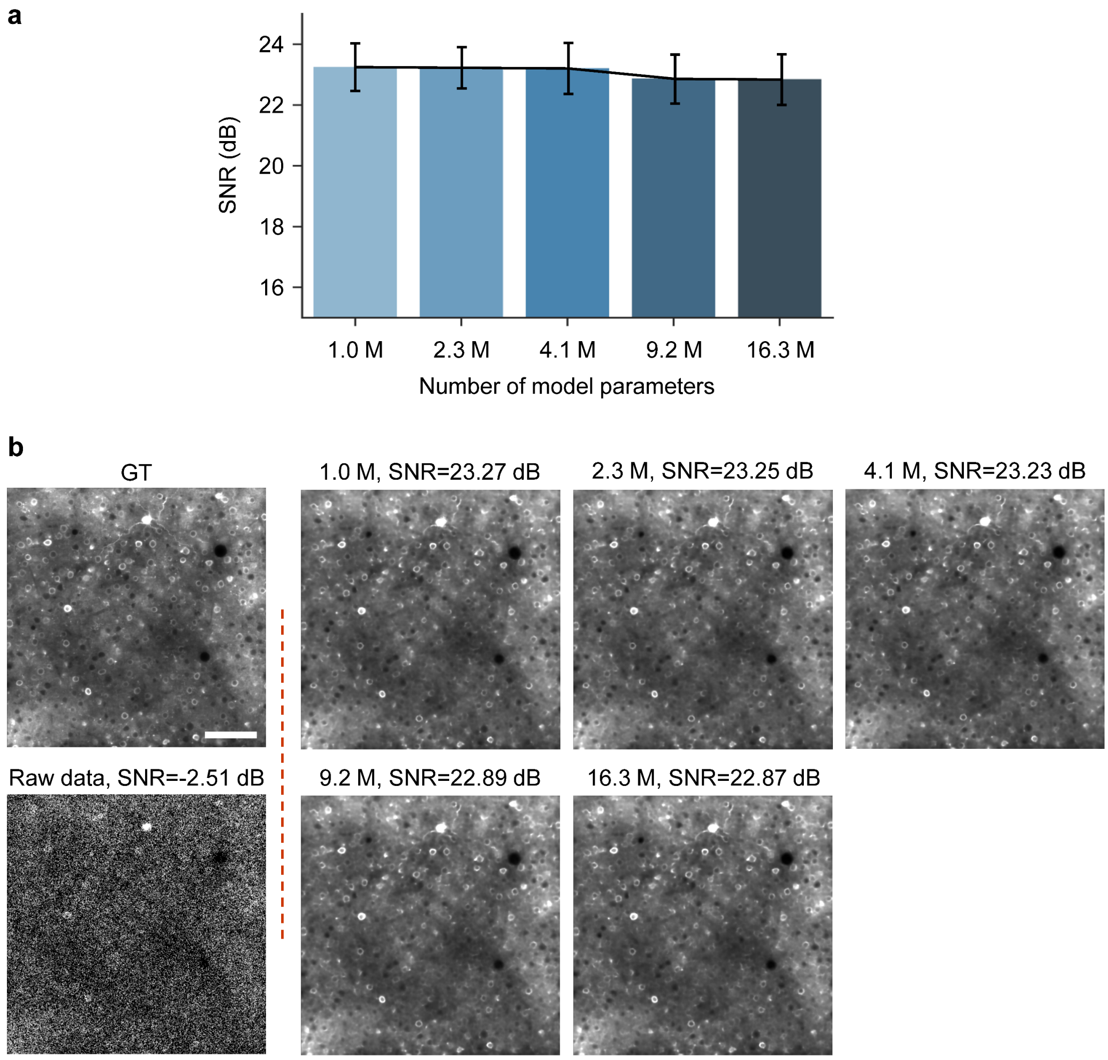


**Supplementary Figure 2**

**Evaluating the performance of different model complexity.**

Simulated calcium imaging data (6000 frames, 30 Hz frame rate, SNR=-2.51 dB) were used in this experiment for quantitative evaluation. The best training epoch was selected by validation. **a**, The relation between denoising performance (output SNR) and the number of model parameters. The bar plot shows mean values and error bars represent the minimum and maximum values. **b**, Example ground truth (GT) images, raw data before denoising, and images after denoising. Scale bar, 100 μm.

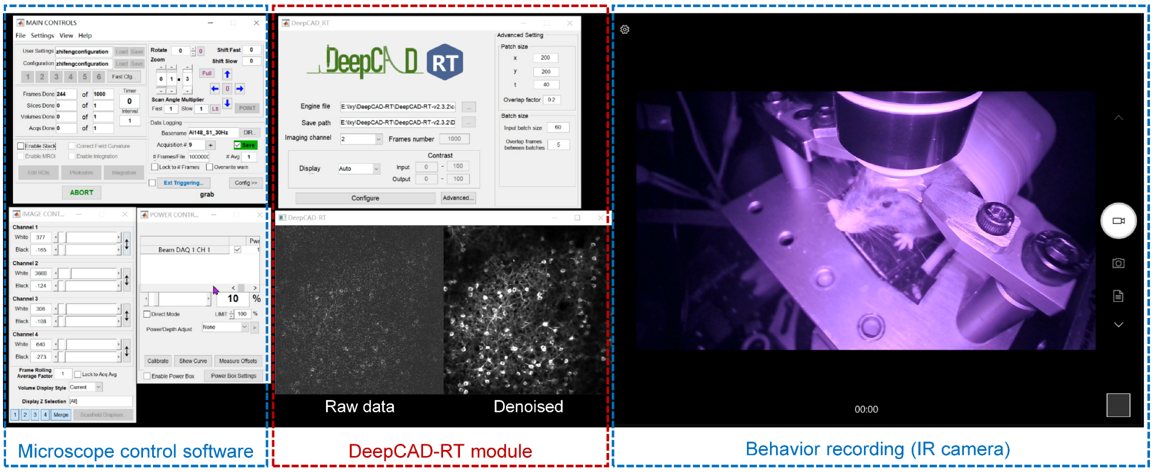


**Supplementary Figure 3**

**Real-time implementation of DeepCAD-RT.**

Real-time denoising was implemented by incorporating DeepCAD-RT into the image acquisition software. Images captured by the microscope were seamlessly fed into DeepCAD-RT, which denoised the input low-SNR images using pre-trained models and displayed denoised images after real-time processing.

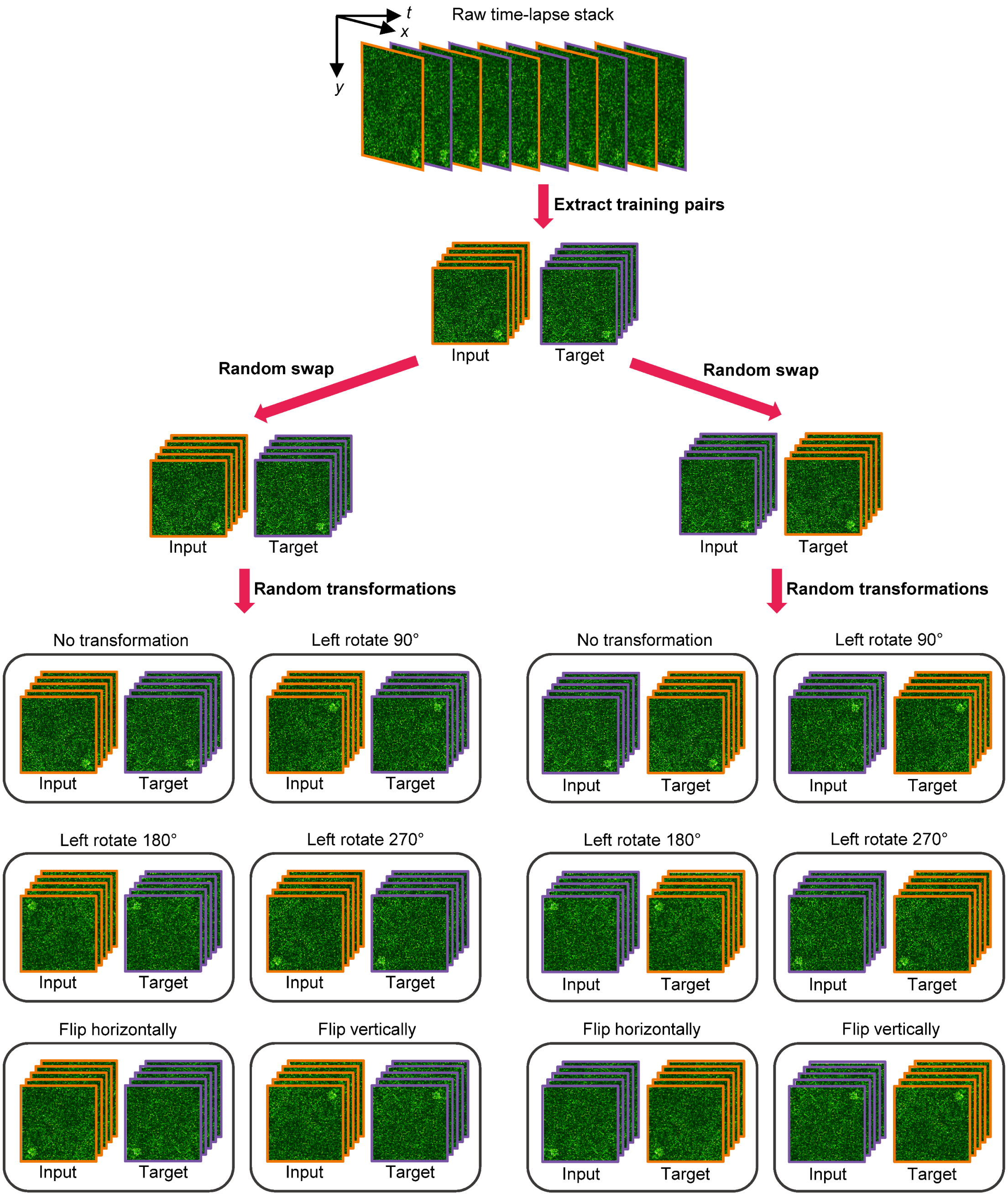


#### Supplementary Figure 4

#### Data augmentation strategy.

Adjacent frames in the original low-SNR stack (xy-t) were divided into two sub-stacks. One as the input volume and the other one as the target volume. Before being fed into the network for training, each training pair was augmented 12-fold through a random swap and six random geometric transformations.

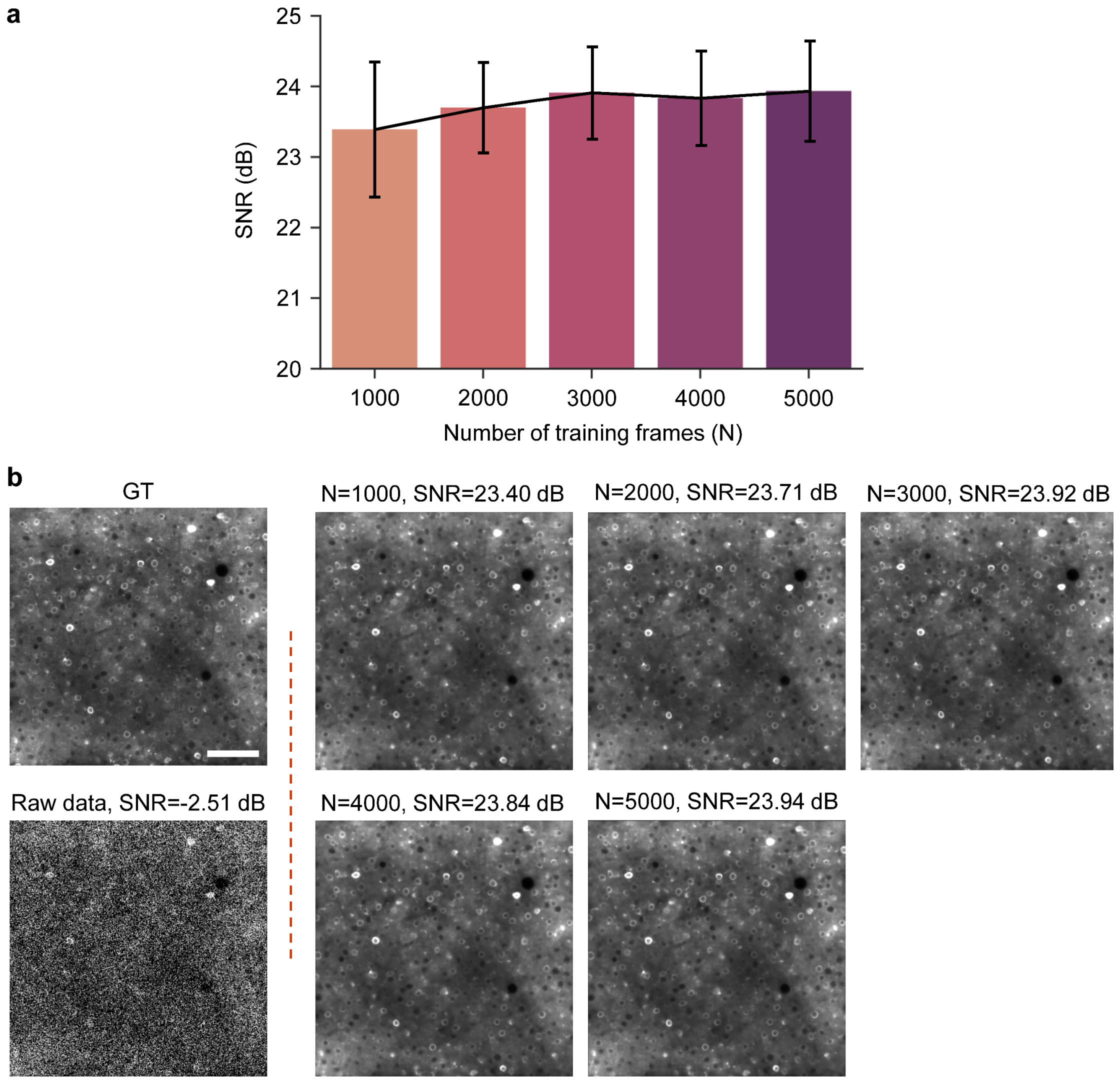


**Supplementary Figure 5**

**Evaluating the data dependency of DeepCAD-RT.**

Simulated calcium imaging data (30 Hz frame rate, SNR=-2.51 dB) were used in this experiment for quantitative evaluation. The network architecture has been simplified (~1.0 million trainable parameters) and data augmentation was applied. Each model was trained for 20 epochs and the last epoch was used for comparison. **a**, The relation between denoising performance (output SNR) and the number of training frames (N). The bar plot shows mean values and error bars represent the minimum and maximum values. **b**, Example ground truth (GT) images, raw data before denoising, and images after denoising. Scale bar, 100 µm.

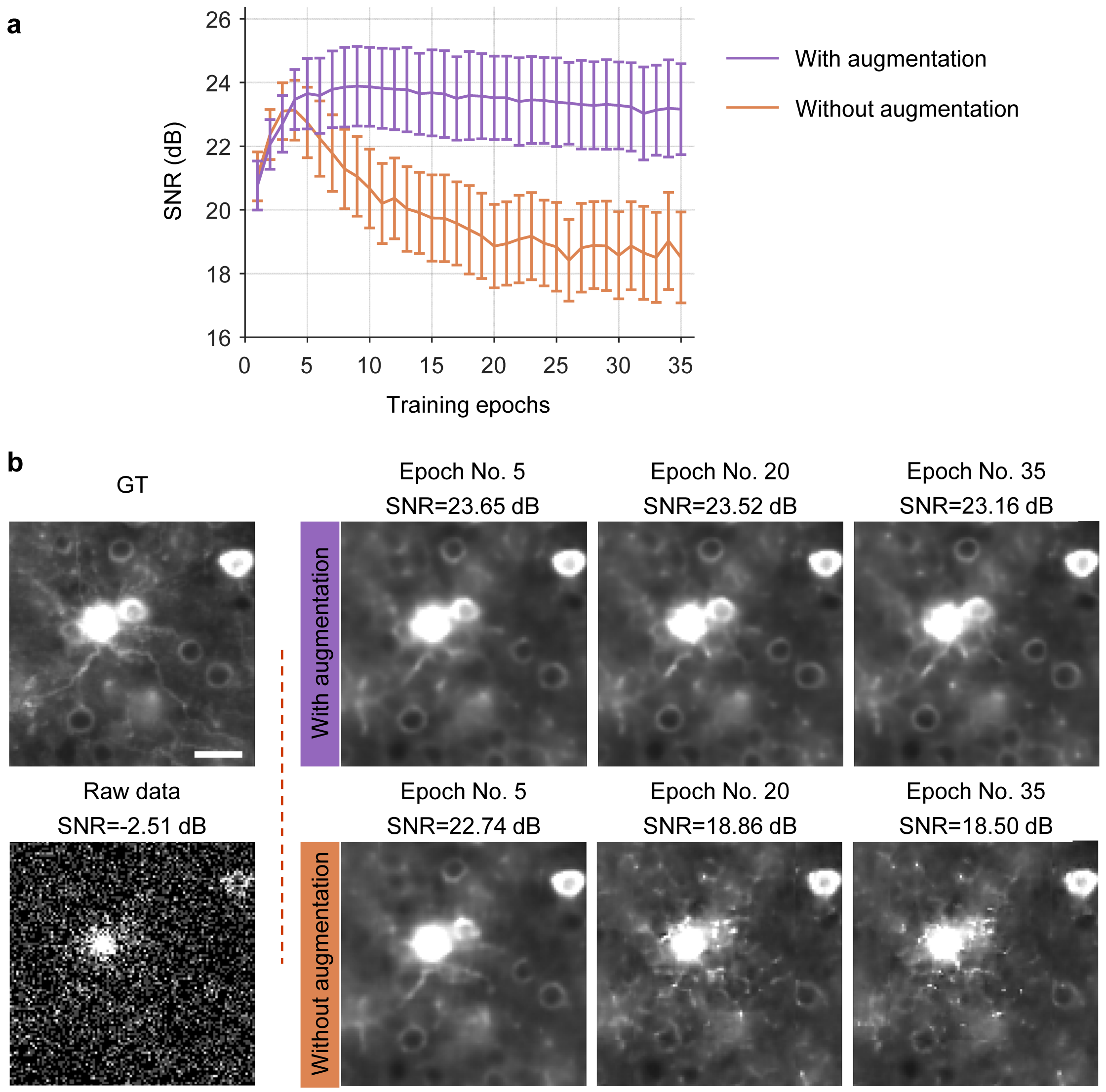


**Supplementary Figure 6**

**Training stability with and without data augmentation method.**

Simulated data (1000 frames, 30 Hz frame rate, SNR=-2.51 dB) were used in this experiment for quantitative evaluation. The network architecture has been simplified (~1.0 million trainable parameters). **a**, Denoising performance (SNR) with the increase of training epoch. Lines represent mean values and error bars represent the minimum and maximum values. **b**, Example ground truth (GT) images, raw data before denoising, and denoising results with and without data augmentation. Scale bar, 20 µm.

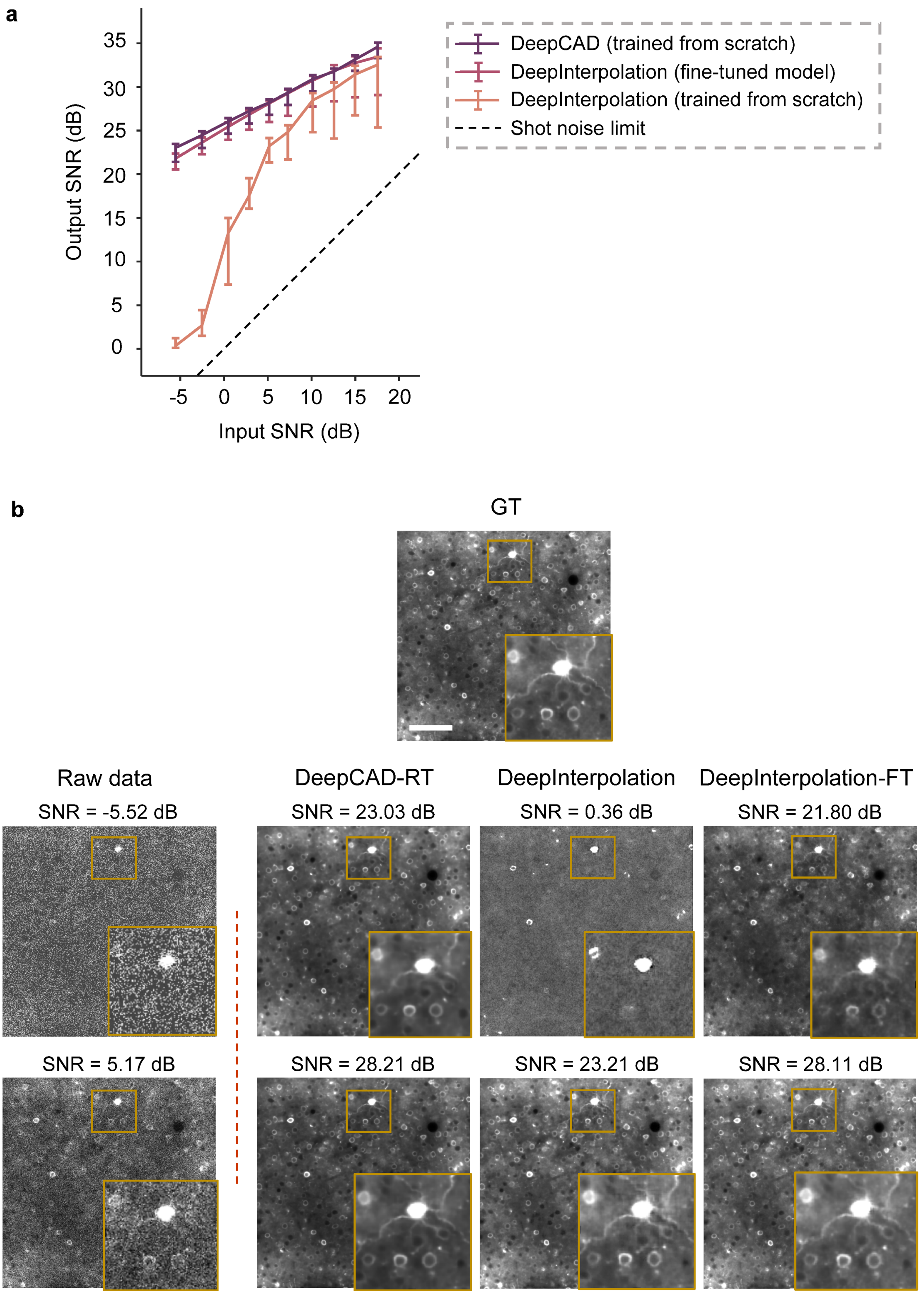

a
Output SNR (dB)
35
30
25
20
15
10
5
0
-5
0
5
10
15
20
Input SNR (dB)
DeepCAD (trained from scratch)
DeepInterpolation (fine-tuned model)
DeepInterpolation (trained from scratch)
Shot noise limit
b
GT
Raw data
DeepCAD-RT
DeepInterpolation
DeepInterpolation-FT
SNR = -5.52 dB
SNR = 23.03 dB
SNR = 0.36 dB
SNR = 21.80 dB
SNR = 5.17 dB
SNR = 28.21 dB
SNR = 23.21 dB
SNR = 28.11 dB

**Supplementary Figure 7**

**Comparing DeepCAD-RT and DeepInterpolation at different noise levels.**

Simulated calcium imaging data (6000 frames, 30 Hz frame rate) were used for the training of all models. DeepInterpolation was implemented with the companion code of relevant papers[4] and two DeepInterpolation models were trained. The first one was trained from scratch. The other model was fine-tuned based on a pre-trained model (pre-trained with 225,000 two-photon images of the Ai93 reporter line) by presenting the training data only once according to the DeepInterpolation paper. DeepCAD-RT (simplified network architecture with ~1.0 million trainable parameters) was trained from scratch for 20 epochs with data augmentation. **a**, The relationship between the input SNR and the output SNR. Lines represent mean values and error bars represent the minimum and maximum values. The black dashed line represents the shot-noise limit (*i.e.*, the input SNR is equal to the output SNR). **b**, Representative images at different input SNRs (-5.52 dB and 5.17 dB). Magnified views of yellow boxed regions are shown at the right bottom of each image. DeepInterpolation, the model trained from scratch. DeepInterpolation-FT, the fine-tuned model. Scale bar, 100 μm.

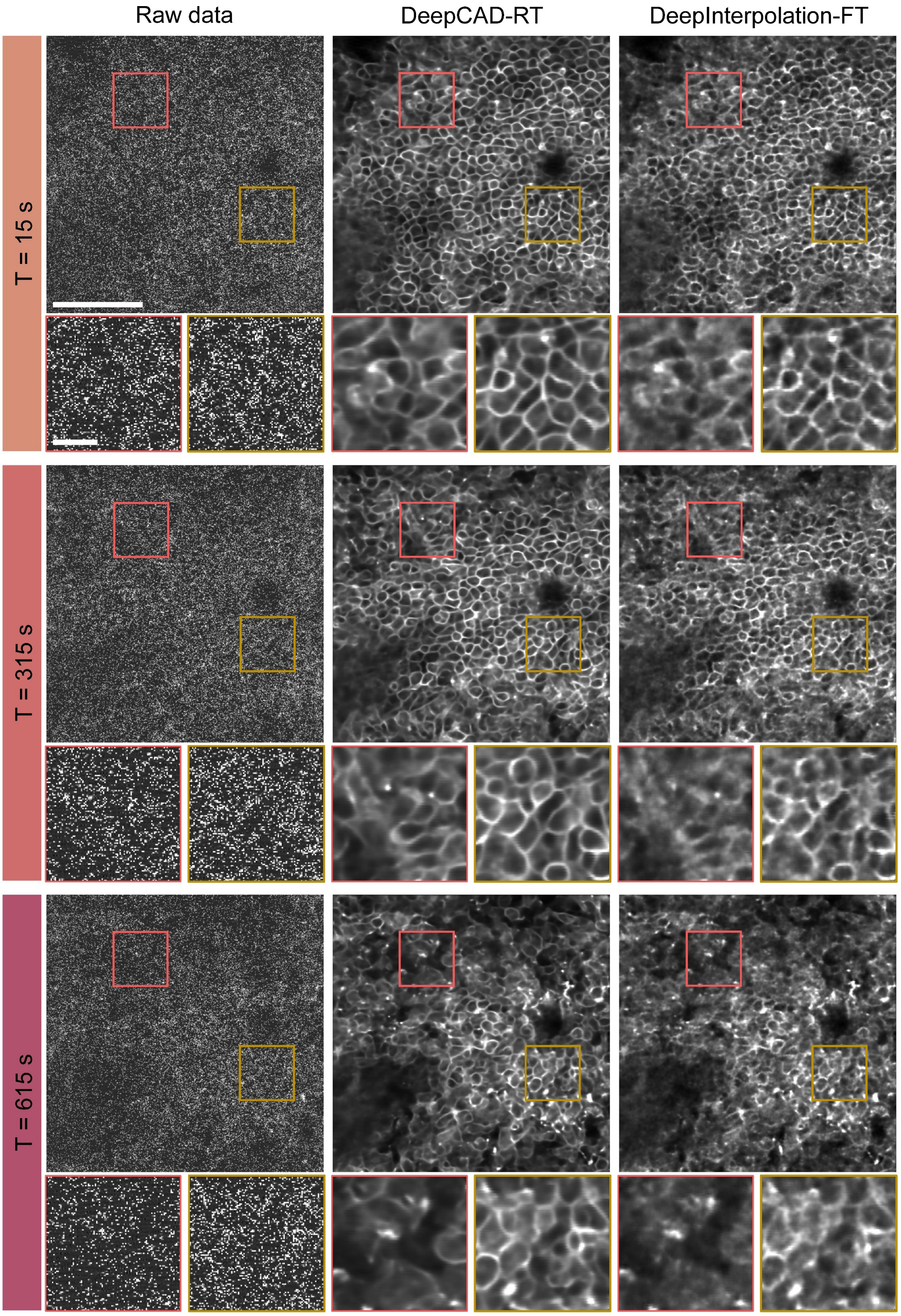
Raw data
DeepCAD-RT
DeepInterpolation-FT
T = 15 s
T = 315 s
T = 615 s

**Supplementary Figure 8**

**Performance comparison between DeepCAD-RT and DeepInterpolation on neutrophil imaging data.**

Neutrophil imaging data (2000 frames, 2 Hz frame rate) were used to train our method and fine-tune DeepInterpolation (DeepInterpolation-FT). Three frames at different moments are demonstrated here and each row shows one frame. Magnified views of boxed regions are shown under each image. Scale bar, 50 μm for the whole field-of-view and 10 μm for magnified views.

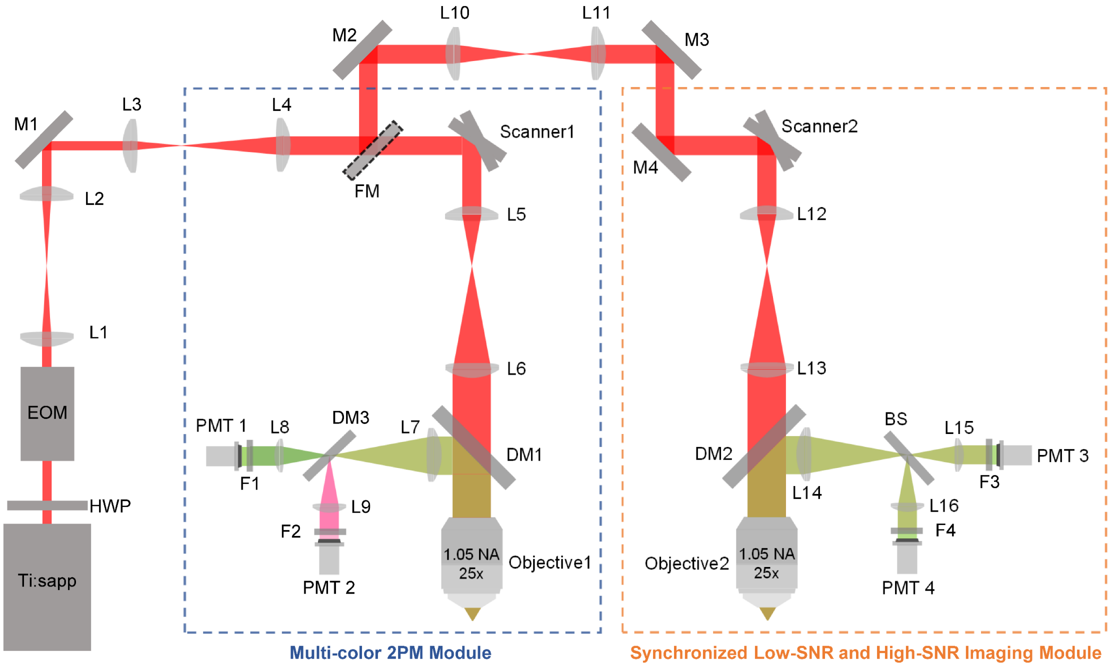


**Supplementary Figure 9**

**Imaging system.**

Our imaging system was composed of a multi-color two-photon module (blue box) and a custom-designed two-photon module to capture synchronized low-SNR and high-SNR images (yellow box). Ti:sapp: titanium-sapphire laser with tunable wavelength; HWP: half-wave plate; EOM: electro-optic modulator; M1-M4: mirrors; L1-L16: lens; Scanner1, Scanner2: galvo-resonant scanners; DM1, DM2: long-pass dichroic mirrors to separate fluorescence signals (green path) from the excitation laser (red path); DM3: short-pass dichroic mirror to separate green fluorescence and red fluorescence. FM: flip mount to alternate between the two modules; F1-F4: emission filters; BS: 1:9 (reflectance: transmission) non-polarizing plate beam splitter; PMT1-PMT4: photomultiplier tubes.

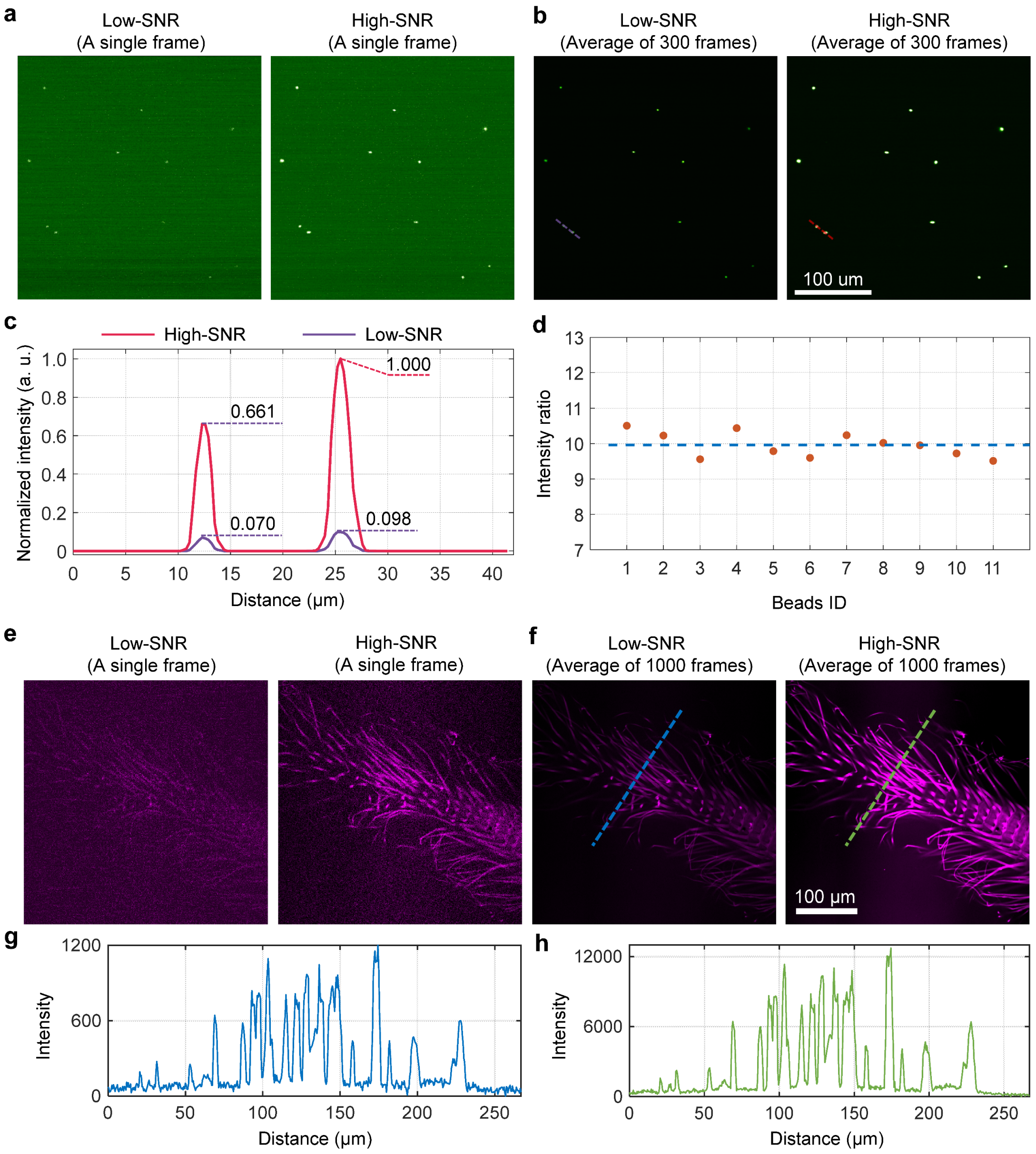


**Supplementary Figure 10**

**System calibration.**

**a**, Example frames captured by the low-SNR detection path (left) and the high-SNR detection path (right). **b**, Average projection of 300 continuously acquired frames. Noise was largely suppressed and underlying fluorescence signals were revealed. **c**, Intensity profiles (normalized to the maximum of high-SNR recording) along the dashed lines in **b**. **d**, The intensity ratios (high-SNR relative to low-SNR) of all 11 fluorescent beads in the FOV. Each point represents one bead and the average intensity ratio is ~10.0 (blue dashed line). **e**, Example images of an insect slice captured by the low-SNR detection path (left) and the high-SNR detection path (right). **f**, Average projection of 1000 consecutive frames. **g, h**, Intensity profiles along the blue and green dashed lines in **f**.

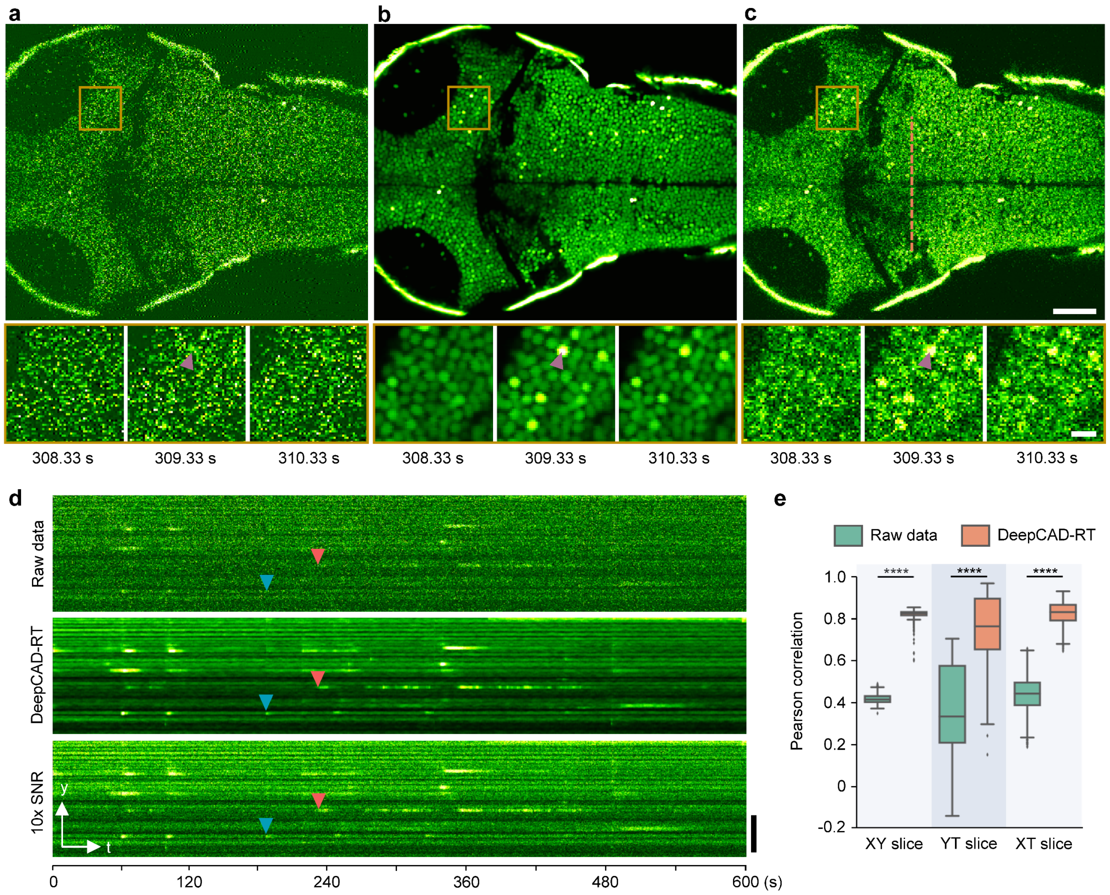


#### Supplementary Figure 11

**Denoising calcium imaging across multiple brain regions in zebrafish.**

**a**, Original low-SNR recording. **b**, DeepCAD-RT enhanced data. **c**, Synchronous high-SNR recording with 10-fold SNR. Magnified views of the yellow boxed region showing calcium dynamics in a 2-second period. Arrowheads point to the same neuron. Scale bar, 50 μm for the large FOV and 10 μm for magnified views. **d**, YT slices along the dashed line in **c**. Two calcium events are indicated with arrowheads of different colors. Scale bar, 50 μm. **e**, Pearson correlation of image slices along all three dimensions before and after denoising. XY slice, N=9000; YT slice, N=400, XT slice, N=485. P values were calculated by one-sided paired t-test. ****$P < 0.0001$ for all comparisons.

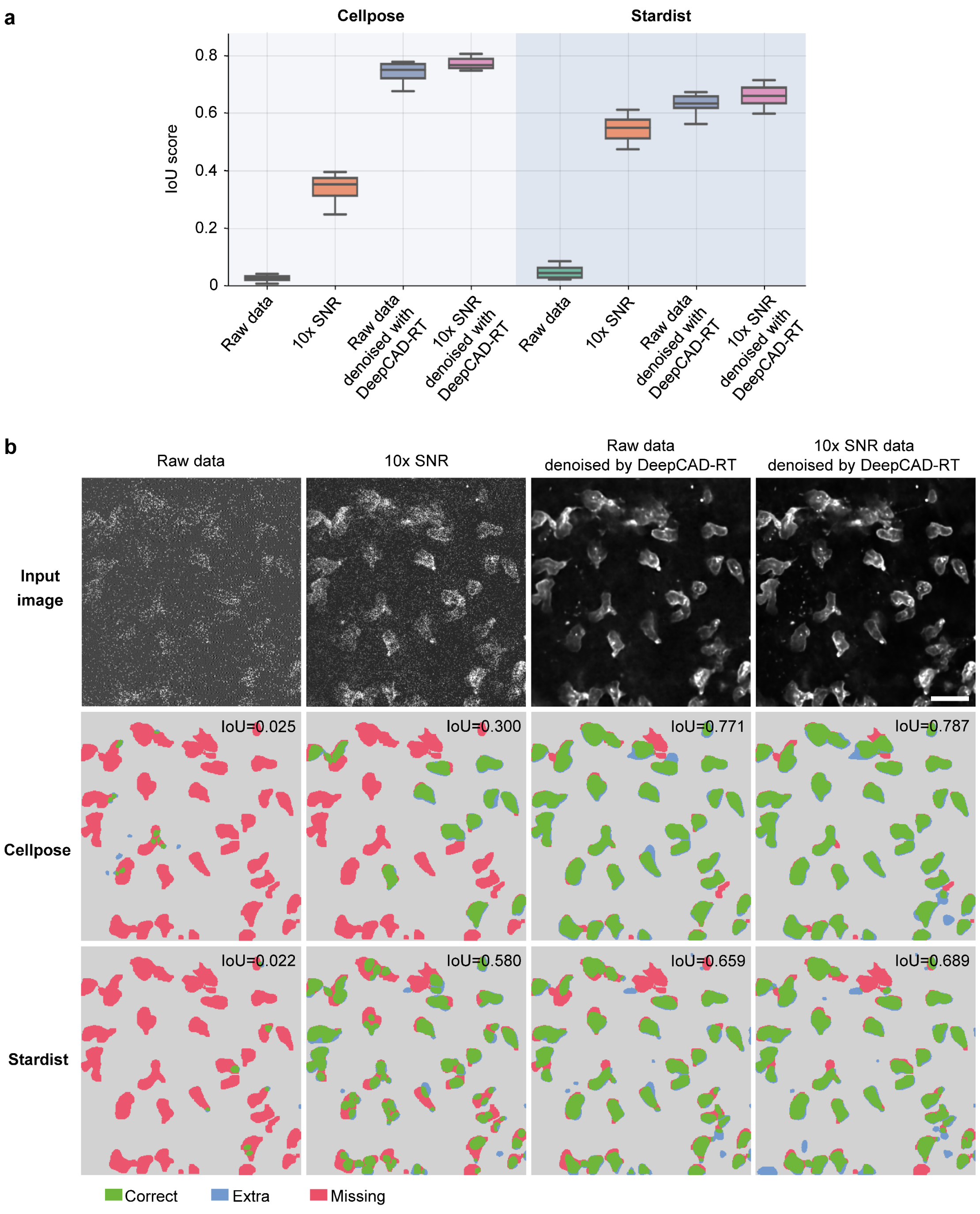


#### Supplementary Figure 12

**The performance of neutrophil segmentation before and after denoising.**

**a**, Segmentation performance of Cellpose[5] and Stardist[6] on raw low-SNR data, synchronous high-SNR data (10× SNR), DeepCAD denoised raw data, and DeepCAD denoised high-SNR data. Intersection-over-Union (IoU) score was used to quantify the segmentation performance. Manually annotated masks were used as the ground truth. **b**, Representative input images and segmented masks. Correctly segmented regions (true positive) are colored green. Missing (false negative) and extra regions (false positive) are colored red and blue, respectively. Scale bar, 20 μm.

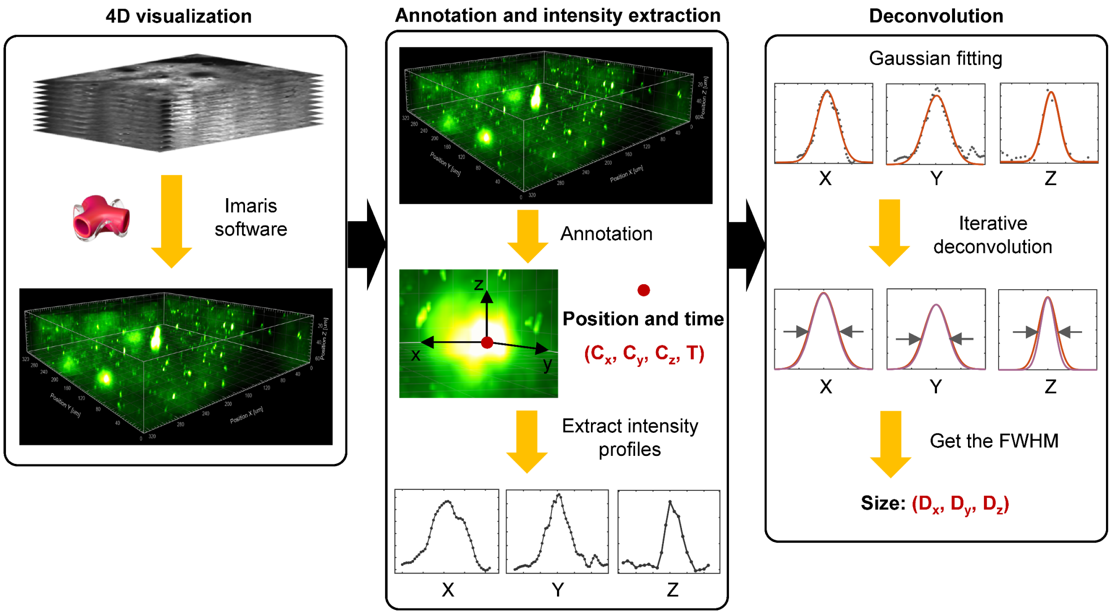


##### Supplementary Figure 13

##### ATP annotation pipeline.

First, low-SNR recordings (4D, xyz-t) were denoised with DeepCAD-RT. Denoised data were visualization in Imaris software. Second, all ATP-release events during the whole imaging session were manually annotated to obtain their position and time. Then, intensity profiles along all three dimensions of each event were extracted from the denoised data. Finally, Gaussian fitting was performed and fitted curves were deconvoluted with the Richardson–Lucy algorithm to eliminate the influence of limited and anisotropic spatial resolution. The full width at half maximum (FWHM) of each deconvoluted curve was extracted as the size of the event.

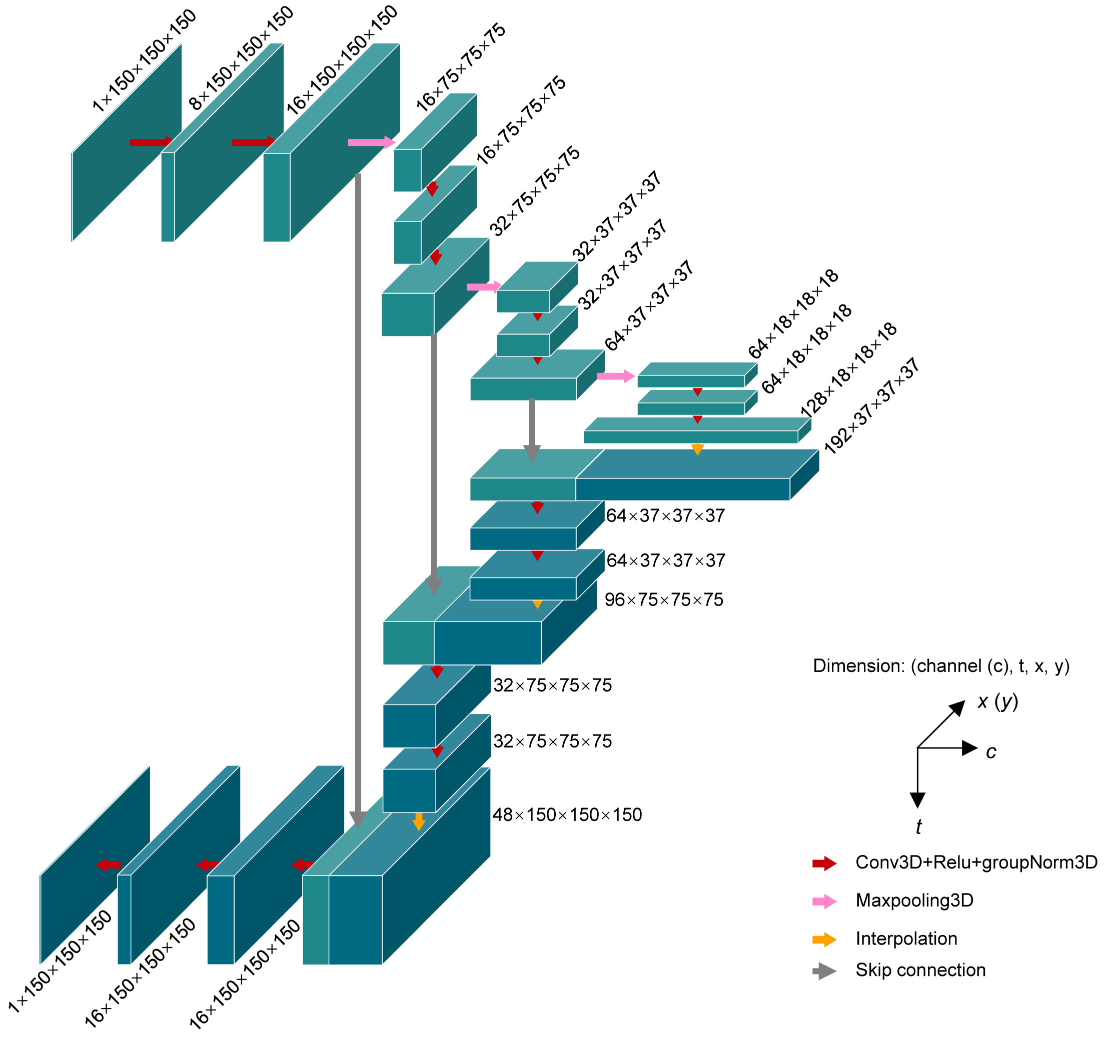


**Supplementary Figure 14**

**Network architecture.**

We used simplified 3D U-net[7] as the network architecture, which is composed of a 3D encoder module, a 3D decoder module, and skip connections from the encoder module to the decoder module. The network architecture was simplified by pruning features in all convolutional layers. The number of trainable parameters was reduced from ~16.3 million (16,315,585) to ~1.0 million (1,020,337) for higher processing speed and less memory consumption.

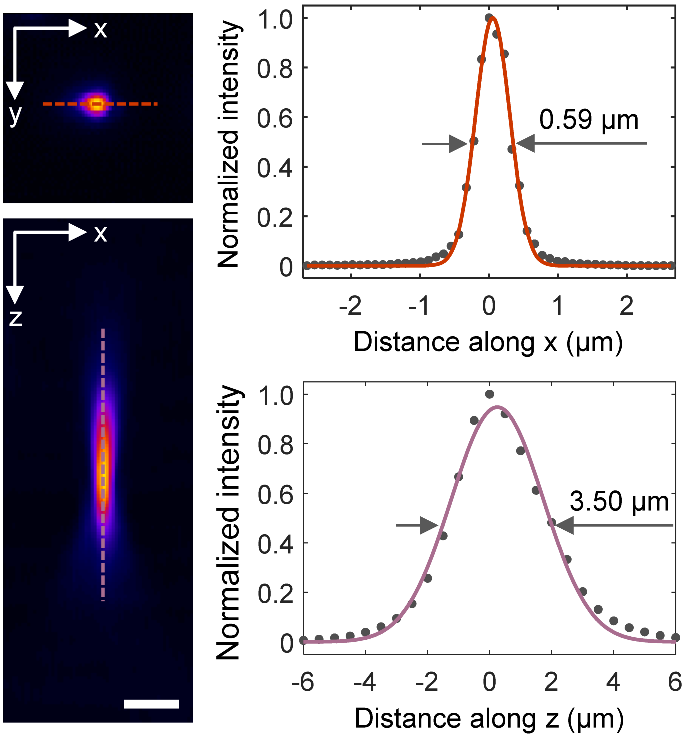


**Supplementary Figure 15**

**System point spread function (PSF).**

The spatial resolution of our imaging system was calibrated by imaging 0.2-μm fluorescent beads. The lateral (XY) and axial (XZ) projections of the system PSF are shown on the left. Normalized intensity profiles of the dashed lines are shown on the right. Solid lines are gaussian fitted curves. The lateral resolution (defined in FWHM) of our system was 0.59 μm and the axial resolution was 3.50 μm. Scale bar, 2 μm.

**Supplementary Table 1**

**Comparison of different model complexity.**

We compared the memory cost, time consumption, and denoising performance of different model complexity using simulated calcium imaging data with clean images for better quantification. All hyper-parameters were kept unchanged except the total number of model parameters. Specifically, all models were trained on 3000 pairs of 150×150×150 (x×y×t) 3D image patches for 20 epochs with one batch size. For inference, a 490×490×300 calcium imaging sequence was split into 75 150×150×150 3D patches and then fed into each model with one batch size. The GPU for training and inference was Nvidia GeForce RTX 3090.

| Model parameters | 16,315,585 (16.3 M) | 9,178,129 (9.2 M) | 4,079,713 (4.1 M) | 2,295,145 (2.3 M) | 1,020,337 (1.0 M) |
|---|---|---|---|---|---|
| Model size (MB)[a] | 62.3 | 35.0 | 15.6 | 8.8 | 3.9 |
| Training memory (GB) | 13.0 | 10.1 | 7.2 | 5.7 | 4.2 |
| Training time (h) | 35.3 | 25.9 | 13.1 | 9.9 | 6.2 |
| Inference memory (GB) | 18.7 | 14.4 | 10.0 | 7.9 | 5.7 |
| Inference time (s)[b] | 53 | 40 | 16 | 12 | 8 |
| Output SNR (dB)[c] | 22.87 | 22.89 | 23.23 | 23.25 | 23.27 |

a. The size of each model file (*.pth).

b. Since the simulated frame rate is 30 Hz, the imaging time for 300 frames is about 10 s.

c. The SNR of the input noisy data is -2.51 dB. The output SNR of each model was measured at the optimal training epoch validated with the ground-truth data.

#### Supplementary Table 2

#### Parameters for the simulation of calcium imaging data

We used NAOMi[1] to generate realistic two-photon calcium imaging data. These simulated noise-free videos were used as the ground truth for comparisons of different denoising methods and evaluations of our method. All simulated data were single-plane time-lapse image sequence. The key simulation parameters are as follows. Those not mentioned in the table all used default values.

| **Physiological parameters** | | **Imaging parameters** | |
|---|---|---|---|
| Sample volume | 500×500×50 μm | FOV[a] | 500×500 μm |
| Average neuron radius | 5.9 μm | Pixel size | 1.02 μm |
| Number of neurons | 1250 | Image size | 490×490 pixels |
| Average firing rate | 0.25 | Frame rate | 30 Hz |
| Vasculature simulation | ON | Imaging depth | 200 μm |
| Background dendrites | ON | Excitation NA[b] | 0.6 |
| Calcium indicator | GCaMP6 | Detection NA | 0.8 |
| Fluorophore concentration | 10 μM | Excitation power | 50 mW |
| **Laser source** | | Number of frames | 6000 |
| Wavelength | 920 nm | Objective focal length | 4.5 mm |
| Repetition frequency | 80 MHz | Brain motion | OFF |
| Pulse width | 150 fs | PSF[c] type | Gaussian |

a. FOV: field-of-view of the microscope.

b. NA: Objective numerical aperture.

c. PSF: point spread function of the optical system.
